# Antagonistic fungal enterotoxins intersect at multiple levels with host innate immune defences

**DOI:** 10.1101/2020.11.23.391201

**Authors:** Xing Zhang, Benjamin Harding, Dina Aggad, Damien Courtine, Jia-Xuan Chen, Nathalie Pujol, Jonathan J. Ewbank

## Abstract

Animals and plants need to defend themselves from pathogen attack. Their defences drive innovation in virulence mechanisms, leading to never-ending cycles of co-evolution in both hosts and pathogens. A full understanding of host immunity therefore requires examination of pathogen virulence strategies. Here, we take advantage of the well-studied innate immune system of *Caenorhabditis elegans* to dissect the action of two virulence factors from its natural fungal pathogen *Drechmeria coniospora*. We show that these two enterotoxins have strikingly different effects when expressed individually in the nematode epidermis. One is able to interfere with diverse aspects of host cell biology, altering vesicle trafficking and preventing the key STAT-like transcription factor STA-2 from activating defensive antimicrobial peptide gene expression. The second, potentially as a consequence of a host surveillance mechanism, increases STA-2 levels in the nucleus, modifies the nucleolus, and causes increased defence gene expression. Our results highlight the remarkably complex and potentially antagonistic mechanisms that come into play in the interaction between co-evolved hosts and pathogens.

## INTRODUCTION

The co-evolution of host and pathogen species can be interpreted as a constant arms race, with rounds of reciprocal adaptations driving diversification and divergence on the molecular and macroscopic scales (Woolhouse et al., 2002). Thus, an examination of the virulence strategies of a host’s natural pathogens can aid the understanding the origin and function of innate immune mechanisms. Our studies focus on the interaction between the nematode *Caenorhabditis elegans* and its natural fungal pathogen *Drechmeria coniospora*. Having dissected in considerable detail *C. elegans* defences, we recently initiated an investigation of *D. coniospora*. *In silico* analysis of the fungal genome revealed a large number of genes and gene families potentially involved in virulence (Lebrigand et al., 2016).

Several approaches exist to address the role of such candidate virulence factors. For example, the corresponding gene could be knocked out in the *D. coniospora* genome, and the effect on virulence measured. Alternatively, the endogenous fungal gene could be engineered so that the resultant protein is tagged to allow its visualisation during infection and/or for the use of biochemistry to identify host protein targets. Both these strategies have been applied to *D. coniospora* (Lebrigand et al., 2016). After these initial successes, the transformation of *D. coniospora* has subsequently proved to be very difficult, if not impossible. In order to study candidate virulence factors, we therefore adopted a strategy to express the corresponding genes directly in *C. elegans*. In a proof of principal, the Falkow laboratory had previously used such a method of heterologous expression, transforming worms with a gene encoding the catalytic subunit of pertussis toxin (PTX) from *Bordetella pertussis*, to find the exotoxin’s conserved target (Darby and Falkow, 2001). More recently, we showed that expressing the *Shigella* virulence factor OspF (a MAP kinase inhibitor) in the epidermis of *C. elegans* blocked the induction of the AMP gene *nlp-29* (Lee et al., 2018), one of the hallmarks of the innate immune response to *D. coniospora* infection (Couillault et al., 2004). Given the toxic nature of some virulence factors, we refined the method to allow tight control of transgene expression, specifically in the adult epidermis of *C. elegans*.

Heat-labile enterotoxins are common and potent virulence factors secreted by pathogenic bacteria. They include cholera and pertussis toxins. They have an αβ_5_ structure, where α is the enzymatically active subunit and the β subunits correspond to the receptor-binding moiety. After the β subunits bind to their specific cell receptor, the toxin is transported into the host cell’s cytosol by endocytosis. The α subunits is then cleaved into α1 and α2 fragments; α1 possesses ADP-ribosylation activity and will modify specific host protein targets (Krueger and Barbieri, 1995). In common with other fungi, *D. coniospora* has no genes encoding enterotoxin β subunits, but does have an unusually large number of genes for enterotoxin α subunit proteins (Wang et al., 2018). The strain used here (derived from ATCC96282) has 19 genes encoding proteins with a “heat-labile enterotoxin alpha chain” (PFAM: PF01375) domain (Lebrigand et al., 2016). Most of them encode proteins with a signal peptide and are expected to be secreted directly into the host cytoplasm, where they could act as toxins, ADP-ribosylating specific targets. The remaining five proteins lack a signal peptide and may represent examples of intracellular toxins with a potential role in defence against nematode predation (see (Kunzler, 2018)). Here, we focused on representative secreted heat-labile enterotoxins and explored the consequences of their expression in *C. elegans* to understand better how *D. coniospora* is able to infect and harm its host.

## METHODS

### *C. elegans* culture

Strains used in this study are listed in the Supplementary Table S1. All strains were maintained on nematode growth media (NGM) and fed *E. coli* strain OP50 (Stiernagle, 2006). Hygromycin-resistant transgenic worms were maintained on OP50-seeded NGM plates containing 0.2 mg/ml hygromycin B (Thermo Fisher). When large populations of aged-matched worms of such strains were required, young adult worms were bleached by standard alkaline hypochlorite treatment (Stiernagle, 2006) and eggs allowed to hatch overnight in 50 mM NaCl with 0.2 mg/ml hygromycin. L1 larvae were washed three times in 50 mM NaCl to remove hygromycin and grown on standard OP50-seeded NGM plates. This proved an efficient way to select for transgenics, while having a minimal effect on worm physiology, as judged by assaying the expression of *irg-1* (Supplementary Figure S1). Otherwise, to perform assays requiring moderate numbers of worms, young adult transgenic worms were transferred from hygromycin-supplemented onto standard NGM plates and their transgenic progeny selected manually on the basis of fluorescent marker gene expression, to avoid the potentially confounding effects of hygromycin (Supplementary Figure S1).

### *D. coniospora* culture and infection

*D. coniospora* (Swe3, derived from ATCC 96282 (Courtine et al., 2020)) spores were amplified by infecting worms every one or two weeks in the lab. The method to grow spores is described in detail in (Powell and Ausubel, 2008). Briefly, sterile 50 mM NaCl was added to plates containing infected worms. A sterile glass microspreader or an L-shaped Pasteur pipette was used to scrape spores gently from the agar surface until the solution became turbid. About 300 μl of the freshly harvested spore solution was then added to a standard 10 cm OP50 plate with 1000-2000 synchronized L4 or young adult worms. The plate was dried in a laminar flow hood and incubated at 25°C for 1 day. Infected worms were harvested with 50 mM NaCl and transferred to an NGM plate supplemented with 100 μg/ml gentamicin and 100 μg/ml ampicillin. The plate was incubated at 25°C for 1 week. Spores were then harvested as above. For assays requiring infected worms, synchronized young adult worms obtained either following treatment with an alkaline hypochlorite solution or using an egg-laying window, were infected by adding 100 μl of a fresh spore solution to a 4 cm OP50 plate (or 200 μl spore solution to 6 cm OP50 plate). Plates were dried briefly under hood and then incubated at 25°C.

### Plasmid construction

*D. coniospora* (Swe3) cDNA was generated as previously described (Lebrigand et al., 2016) and purified by QIAquick PCR Purification Kit (Qiagen, Cat No./ID: 28104). The fungal gene RJ55_04834, without the 5’ sequence corresponding to the predicted signal peptide, with an engineered ATG, and without the stop codon, was amplified from cDNA by PCR with Gibson assembly 5’ and 3’ primers that included *NotI* and *ClaI* sites, respectively (Supplementary Table S2). FLAG and tev sequences were synthesised by Integrated DNA Technologies (Leuven, Belgium). mKate2 was amplified from the plasmid pNP152 (Sinner et al., in press) with Gibson assembly primers. The vector backbone was amplified by PCR using the destination vector pSX103 that contains the promoter of *col-19* (Xu and Chisholm, 2011), a generous gift from Andrew Chisholm, as the template. All the fragments were assembled using the Gibson assembly protocol (Gibson et al., 2009) to give pZX12. The degron sequence was amplified from DNA extracted from the worm strain PX627 (Kasimatis et al., 2018) and inserted into *AscI*-digested pZX12 using Gibson assembly to give pZX17. To make other plasmids, pZX17 was double digested by *NotI* and *ClaI*, and the RJ55_04834 gene fragment replaced with the appropriate alternative *D. coniospora* gene fragment, generated as described above. *rps-0p*::hygR was amplified from pSO5.3 (Lee et al., 2018) and inserted into the pSX103 vector (double digested with *KpnI* and *NarI*) using Gibson assembly to give the pZX13 plasmid.

### Microinjection

Microinjections were performed using 20 ng/μl of the virulence factor construct, 20 ng/μl *rps-0p*::hygR and coinjection marker *unc-122p*::GFP, a kind gift from Jean-Louis Bessereau, at a concentration of 40 ng/μl into JDW141 (*eft-3p*::TIR::P2A:::BFP-NLS-degron::*tbb-2* 3'UTR). This strain, for use with the auxin-inducible degron system, with an internal degradation control (Ashley et al., 2020), was a generous gift of Jordan Ward. To generate the control hygR;*fr*Is7 strain (IG1864), 60 ng/μl *rps-0p*::hygR and 60 ng/μl *unc-122p*::GFP were microinjected into IG274 (+;*frIs7*) worms (Pujol et al., 2008a).

### RNA extraction, reverse transcription and quantitative PCR

Worms were harvested and washed three times with 50 mM NaCl and pelleted by centrifugation before Trizol extraction (Thermo Fisher Scientific) following the manufacturer’s instructions. Reverse transcription was performed using High-Capacity cDNA Reverse Transcription Kit (Invitrogen). Quantitative real-time PCR was performed as described (Pujol et al., 2008b) by using SYBR™ Green PCR Master Mix (TaKaRa). Values were normalized to those of *act-1* and were analyzed by the cycling threshold method using the appropriate qRT-PCR primers (Supplementary Table S2). Control and experimental conditions were tested in the same run.

### Microscopy and image analysis

Worms were picked into a drop of 5 mM levamisole on a 2% agarose pad on a glass slide and observed using a Leica DMRBE microscope. Fluorescent images were taken with a Zeiss AxioCam HR digital colour camera and Axio-Vision Rel. 4.6 software (Carl Zeiss AG). Confocal microscopy used a Zeiss LSM 780. All image processing was done using Fiji software (Schindelin et al., 2012). Comparative quantifications were performed with image sets acquired on same day. The area and Feret’s (caliper) diameter for lateral hyp7 nucleoli were measured with an automatic particle analysis method. Feret’s diameter represents the longest distance between any two points along an object’s boundary. For better information extraction, minimizing the background noise and to avoid over and under estimation, automatic thresholding was applied to images of each nucleolus, selected after pseudo-colouring based on pixel intensity and smoothing to discriminate better the area of interest. For any data that did not have a normal distribution (determined with a Shapiro-Wilk test), statistical significance was determined using a nonparametric Mann Whitney test (GraphPad Prism software).

### Analyses with the Biosort worm sorter

Fluorescent protein expression of reporter strains was quantified with the COPAS (Complex Object Parametric Analyzer and Sorter) Biosort system (Union Biometrica; Holliston, MA) as described (Pujol et al., 2008a). For each strain, a minimum of 150 synchronized young adult worms were analyzed for length (assessed as TOF, time of flight), optical density (assessed as extinction) and Green and/or Red fluorescence (GFP/Red). Raw data were filtered on the TOF for adult worms (typically 300 ≤ TOF ≤ 1500). Statistical significance was determined using a non-parametric analysis of variance with a Dunn’s test (GraphPad Prism).

### RNA interference

RNAi bacterial clones were obtained from the Ahringer or Vidal libraries (Kamath and Ahringer, 2003; Rual et al., 2004). RNAi bacteria were seeded on NGM plates with the appropriate antibiotics. Worms were transferred onto RNAi plates as L1 larvae and cultured at 20°C or 25°C as indicated.

### Lifespan

L4 worms were manually transferred to small (4 cm) plates containing NGM agar seeded with *E. coli* OP50. Typically, 10 worms were put into each of 5 plates per strain. Worms were grown at 25°C and the surviving and dead worms were counted every day. Worms were transferred to new plates every day at the start of the experiment to eliminate the larvae of the next generation. Once worms had stopped producing viable eggs, they were kept on the same plates and the worms that no longer responded to light touch were picked out as scored as dead.

### Survival upon *D. coniospora* infection

L4 or young adult worms were manually transferred to 4 cm plates containing NGM agar seeded with *E. coli* OP50. Fresh spores were harvested and spread on the plate. Normally, 1 × 10^8^ spores were used for infecting about 100 worms on a 4 cm OP50 plate. After infection (either 8 hours or overnight), for each experimental condition, 25 worms were put into each of 4 wells of a 12-well plate containing NGM agar seeded with *E. coli* OP50. Images of each well were collected automatically every 24 minutes using a custom system that will be described elsewhere. The images were examined, and worms scored as dead when they no longer exhibited movement between successive images.

### Cuticle fragility test

The cuticle fragility was tested by measuring the time to cuticle rupture in bleach as previously described (Tong et al., 2009).

### Cycloheximide (CHX) treatment

CHX (40 mM in DMSO) was added to OP50-seeded NGM agar plates to a final concentration of 500 μg/ml and dried in a laminar flow hood and then stored for at least 1 day. Young adult worms that had been grown at 20°C on OP50 NGM plates were transferred to the plates with CHX for 6h.

### Preparation of worms for biochemistry

Large quantities of worms were prepared using enriched NGM agar medium (NGM+; 3 g NaCl, 20 g peptone, 25 g agar, 1 ml of 5 mg/ml cholesterol (in ethanol) in 975 ml of H_2_0 autoclaved, cooled then supplemented with 1 ml of 1 M CaCl_2_, 1 ml of 1 M MgSO_4_, 25 ml of 1 M Phosphate buffer, 1 mL of 100 mg/mL ampicillin), seeded with 10x concentrated HT115 *sta-1*(RNAi) clone, and grown for 20-24 h at 37°C to obtain a thick bacterial lawn. During the amplification of strains, to select transgenic worms, seeded plates were supplemented with hygromycin at a final concentration of 0.2 mg/ml. In the case of worms expressing DcEntA and DcEntB, plates were additionally supplemented with auxin at a final concentration of 1 mM to limit any potential toxicity due to expression of the virulence factors.

Mixed stage worm populations were collected by using 50 mM NaCl 0.05% Triton X-100 in 15 ml tubes, then washed three times in 50 mM NaCl prior to standard alkaline hypochlorite treatment. The recovered eggs were then washed three times and allowed to hatch overnight in 3 ml of 50 mM NaCl supplemented with 0.2 mg/ml hygromycin, with gentle agitation. The synchronised L1 worms were then added to NGM + plates, and grown at 25°C until they reached the young adult stage. The expression of the chimeric virulence protein was confirmed by the observation of the expected red fluorescence signal using a dissecting fluorescence microscope. Worm samples were then collected by using 50 mM NaCl, 0.05% Triton X-100 in 15 ml tubes and then washed 3-4 times in 50 mM NaCl, until the supernatant was cleared of bacteria, before freezing the worm pellets at −80°C.

### Immunoprecipitation assay

Pellets of synchronised young adult worms (0.8-1 ml) were thawed in the presence of an equal volume of 2x lysis buffer (75 mM Hepes, pH 7.5; 1.5 mM EGTA; 1.5 mM MgCl_2_; 150 mM KCl; 15% glycerol; 0.075% NP-40 to which Roche cOmplete™ Mini tablets containing a protease inhibitor cocktail were added just prior to use). Thawed worm pellets were then flash frozen in liquid nitrogen and then crushed using a pestle and mortar on dry ice prior to sonication in Diagenode 15 ml tubes (C30010017) using a Diagenode BioRuptor Pico (10 cycles 15 s on; 45 s off) cooled to 4°C. Lysates were then transferred to 2 ml Eppendorf tubes, the concentration of NP-40 adjusted to 0.5% and centrifuged for 10 min at 8000 rpm at 4°C. The supernatant was then transferred in 900 μl aliquots to fresh Eppendorf tubes and incubated with gentle head-over-tail agitation for 2 h at 4°C with 30 μl of anti-RFP or control beads (Chromotek rtma-2 and bmab-20, respectively) that had been washed in 1x lysis buffer.

### Protein in-gel digestion

Proteins were separated briefly in a 4-12% NuPAGE Bis-Tris gel, stained with coomassie blue and cut into small gel cubes, followed by destaining in 50% ethanol/25 mM ammonium bicarbonate. The proteins were then reduced in 10 mM DTT at 56 °C and alkylated by 50 mM iodoacetamide in the dark at room temperature. Afterwards, proteins were digested by trypsin (1 μg per sample) overnight at 37 °C. Following peptide extraction through sequential incubation of gel cubes in 30% and 100% acetonitrile, the sample volume was reduced in a centrifugal evaporator (Eppendorf) to remove residual acetonitrile. The resultant peptide solution was purified by solid phase extraction in C18 StageTips (Rappsilber et al., 2003).

### Liquid chromatography tandem mass spectrometry

Peptides were separated in an in-house packed 50 cm analytical column (inner diameter: 75 μm; ReproSil-Pur 120 C18-AQ 1.9 μm resin, Dr. Maisch GmbH) by online reversed phase chromatography through a 90 min gradient of 2.4-32% acetonitrile with 0.1% formic acid at a nanoflow rate of 250 nl/min. The eluted peptides were sprayed directly by electrospray ionization into an Orbitrap Exploris 480 mass spectrometer (Thermo Scientific). Mass spectrometry measurement was conducted in data-dependent acquisition mode using a top15 method with one full scan (resolution: 60,000, target value: 3 × 10^6^, maximum injection time: 28 ms) followed by 15 fragmentation scans via higher energy collision dissociation (HCD; normalised collision energy: 30“, resolution: 15,000, target value: 1 × 10^5^, maximum injection time: 40 ms, isolation window: 1.4 m/z). Precursor ions of unassigned, +1, +7 or higher charge state were rejected for fragmentation scans. Additionally, precursor ions already isolated for fragmentation were dynamically excluded for 25 s.

### Mass spectrometry data analysis

Raw data files were processed by MaxQuant software package (version 1.6.5.0) (Cox and Mann, 2008) using Andromeda search engine (Cox et al., 2011). Spectral data were searched against a target-decoy database consisting of the forward and reverse sequences of WormPep release WS275 (28,466 entries), UniProt *E. coli* K-12 proteome release 2020_01 (4,403 entries), the corresponding transgenic fusion protein and a list of 246 common contaminants. Trypsin/P specificity was selected. Carbamidomethylation of cysteine was chosen as fixed modification. Oxidation of methionine and acetylation of the protein N-terminus were set as variable modifications. A maximum of 2 missed cleavages were allowed. The minimum peptide length was set to be 7 amino acids. At least one unique peptide was required for each protein group. False discovery rate (FDR) was set to 1% for both peptide and protein identifications. A separate database search was performed in strain DcEntA to identify mono-ADP-ribosylation (MAR; C_15_H_21_N_5_O_13_P_2_, *m/z* 541.0611) sites on residues CDEKNRST as variable modifications and ribose, adenosine, AMP and ADP as diagnostic ions (Martello et al., 2016).

Protein quantification was performed using the LFQ label-free quantification algorithm (Cox et al., 2014). Minimum LFQ ratio count was set to one. Both the unique and razor peptides were used for protein quantification. The “match between runs” option was used for transferring identifications between measurement runs allowing a maximal retention time window of 0.7 min. All mass spectrometry raw data have been deposited to the PRIDE repository (Perez-Riverol et al., 2019) with the dataset identifier PXD021929. DcEntA and DcEntB are referenced as pZX26 and pZX25, respectively. This dataset also includes unpublished pull-down data for three other structurally-unrelated virulence factors (Harding *et al*., manuscript in preparation) used to identify proteins that bound in a non-specific manner, as described below.

Statistical data analysis was performed using R statistical software. Only proteins quantified in at least two out of the three RFP pull-down replicates were included in the analysis. LFQ intensities were log-transformed. Imputation for missing values was performed for each pull-down replicate by random picking from a normal distribution that simulated low intensity values below the noise level. The LFQ abundance ratio was then calculated for each protein between the RFP pull-downs and the controls. Significance of the enrichment was measured by an independent-sample Student’s *t* test assuming equal variances. Specific interaction partners were then determined in a volcano plot where a combined threshold (hyperbolic curve) was set based on a modified *t*-statistic (SAM, significance analysis of microarrays) (Li, 2012; Tusher et al., 2001). Proteins that passed the combined threshold in a second strain (PRIDE repository PXD021929) were considered as potential cross-reactive, non-specific binders and were filtered out from the volcano plot and summary spreadsheets.

### *In silico* analyses

BLASTP searches were conducted at the NCBI on 09/11/20, using default parameters, but excluding *D. coniospora* entries. We used the command-line version of NoD (v1.3b) (Scott et al., 2011) available as a Conda package (https://anaconda.org/bioconda/clinod). In addition to the default parameters, we used “-clean_sequence” as some *D. coniospora* proteins in the dataset (Lebrigand et al., 2016) contain Xs in their sequence. The default output of NoD was parsed using a Python script (enclosed as a Jupyter notebook; available on request). Gene class enrichment analyses used Wormbase release WS277.

## RESULTS

### Heterologous expression of a single *D. coniospora* protein can be lethal to *C. elegans*

As part of our investigation of the virulence mechanisms of *D. coniospora*, we adopted the strategy of expressing candidate fungal proteins in a controlled manner through transgenesis in *C. elegans*. For endoparasites like *D. coniospora*, virulence factors are often secreted into the host. More than 600 proteins from *D. coniospora* are predicted to be secreted. Among them, 225 previously lacked identifiable protein domains or homologues in other species (Lebrigand et al., 2016). BLASTP searches revealed similar proteins in Genbank (identity >28“; e value < 10^−5^) for 126 of them, and 20 with borderline hits. The remaining 79 are still *D. coniospora* specific proteins of completely unknown function (Supplementary Table S3).

Our initial efforts focused on RJ55_02698 (previously referred to as g2698 (Lebrigand et al., 2016)), a protein similar to Abrin-a from the nematode-trapping fungus *Drechslerella brochopaga*. Abrins are highly toxic as they contain a “ribosome inactivating protein” domain (RIP; PFAM: PF00161). RIPs exhibit RNA N-glycosidase activity and depurinate the 28S rRNA of the eukaryotic 60S ribosomal subunit, thereby inhibiting protein synthesis (Walsh et al., 2013). Despite the fact that we expressed RJ55_02698 under the control of a promoter (*col-19*) that is active principally in the adult epidermis (Cox and Hirsh, 1985; Liu et al., 1995), we had difficulty obtaining many adult transgenic worms. Most transgenic worms were visibly sick and died during development. As the *col-19* promoter has some activity during development, peaking at the L2 stage at a level 50-fold lower than in young adults (Grun et al., 2014) (Supplementary Figure S2A), this morbidity was probably due to the high toxicity of even small quantities of the fungal protein. Before dying, the larvae and rare adults exhibited a wide range of different phenotypes, with variable penetrance, including defects in moulting, rolling, egg retention, and multiple vulvae (Supplementary Figure S2B-F). This presumably reflected stochastic loss of protein expression in the epidermis.

To avoid the potential deleterious effect of virulence factor expression during development, we applied a dual control strategy to limit fungal protein production. In addition to knocking down any gene expression by growing worms on bacteria expressing dsRNA targeting the corresponding sequence, we engineered the proteins as chimeric constructs, including an auxin-inducible degron (Zhang et al., 2015). We also included tags for microscopy and biochemistry (mKate2 and Flag, respectively; Figure 1A). This allowed us to generate stable transgenic lines of *C. elegans* that only expressed the candidate fungal virulence factors when the RNAi and/or auxin-dependent inhibition was removed.

**Figure 1.**
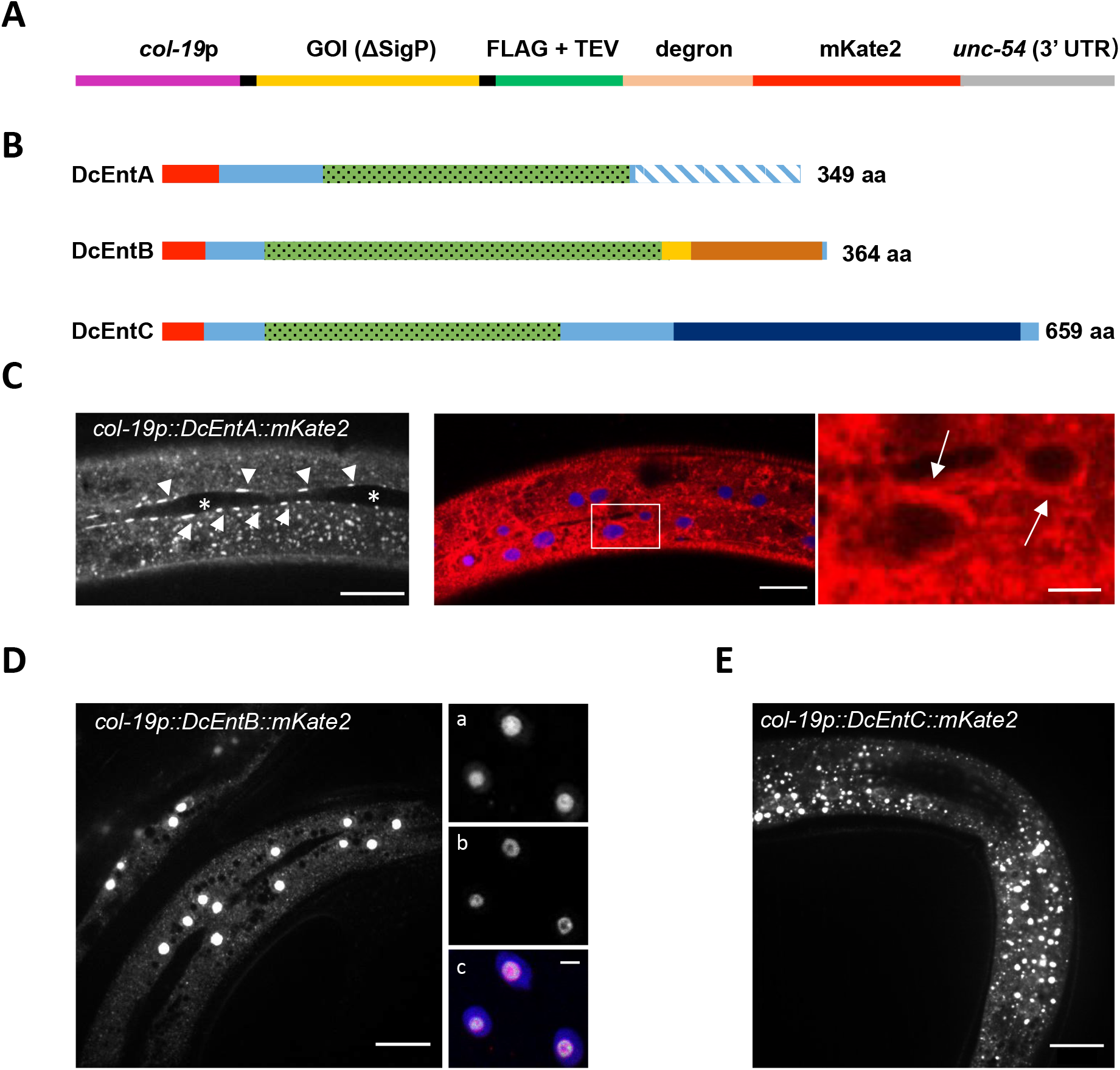
Three enterotoxins have different expression patterns in the epidermis. (**A**) Schematic overview of the plasmid insert used to express virulence factors. The candidate gene of interest (GOI), corresponding to a virulence factor without its signal peptide (ΔSigP) and stop codon was cloned between the *col-19* promoter and *unc-54* 3’ UTR, and expressed as a fusion protein with FLAG, tobacco etch virus (TEV) protease cleavage site, degron and mKate2. (**B**) Schematic overview of three selected enterotoxins from *D. coniospora*. The signal peptide is represented in red, the heat-labile enterotoxin alpha chain domain (PFAM: PF01375) in green, a *D. coniospora* specific 90-residue C-terminal domain in DcEntA (hatched blue), the DcEntB nucleolar targeting sequence predicted by NoD (Scott et al., 2011) in yellow, the regions similar (p=9.44e^−3^) to part of an adenylate kinase domain (PRK13808), in brown, and DNA double-strand break repair ATPase Rad50 (PRK03918; 4.4e^−5^) in dark blue for DcEntC. Representative confocal fluorescence images of young adult worm expressing DcEntA::mKate2 (IG1926; **C**), DcEntB::mKate2 (IG1925; **D**), and DcEntC::mKate2 (IG1880; **E**). (**C**) Left panel: DcEntA (arrowheads) adjacent to seam cells (asterisks). Middle: DcEntA (red) accumulation around nuclei (blue) is highlighted with arrows in the right panel showing a magnified view of the boxed area. Scale bars, left and middle, 20 μm, right, 5 μm. (**D**) Left panel: young adult IG1925 worm expressing DcEntB::mKate2. Scale bar, 20 μm. Right: Image of young adult IG1984 worm expressing DcEntB::mKate2, FIB-1::GFP, and TIR::BFP-NLS (a: red channel; b: green channel; c: overlay red, green and blue channels). Scale bar, 5 μm. (**E**) DcEntC::mKate2 appears as a punctate cytoplasmic pattern. Scale bar, 20 μm.

### Enterotoxins have distinct expression patterns and can make worms sick when expressed in the epidermis

We used this modified approach to address the function of representative members of the large family of *D. coniospora* enterotoxin α (PF01375) domain proteins. We selected three candidates (RJ55_08534/g7949, RJ55_03010/g2819 and RJ55_07324/g6833) that we refer to here as DcEntA, DcEntB and DcEntC, respectively (Figure 1B). When expressed as fusion proteins, each gave a distinct pattern of intracellular localisation in the hyp7 syncytium. DcEntA was enriched in the perinuclear region as well as at the cell membrane, adjacent to seam cells (Figure 1C). DcEntB appeared to be restricted to the nucleolus, consistent with the presence of a predicted nucleolus-localisation signal (NoLS) in its primary sequence (Figure 1B, D). This was confirmed by its co-localisation with the known nucleolar marker FIB-1::GFP (Yi et al., 2015) (Figure 1D). DcEntC, on the other hand, gave a punctate, cytoplasmic pattern (Figure 1E). Any strategy of expressing chimeric proteins under the control of a heterologous promoter runs the risk of experimental artefacts. The very distinct patterns seen for the 3 chimeric DcEnt proteins, however, suggests that their localisation was not unduly influenced by the added non-fungal sequences that are common to all 3 proteins.

The expression of DcEntA and DcEntB provoked very visible phenotypes. The worms appeared sick and were short-lived under standard culture conditions. Worms expressing DcEntC, however, had subtle phenotypes, an almost normal morphology and behaviour, and had a life-span that was indistinguishable from control animals (Figure 2A, B; Supplementary Table S4). In all cases, we were able to maintain viable strains under standard culture conditions. In subsequent experiments, we therefore conducted experiments on worms that had not been exposed to auxin, avoiding its possible confounding side-effects (Bhoi et al., 2020), and concentrated principally on the characterisation of DcEntA and DcEntB. Upon normal handling, adult DcEntA-expressing worms, but not DcEntB-expressing worms, appeared more prone to break apart. This was confirmed in the standard test of cuticle fragility, wherein their time to rupture was significantly shorter than control worms (Figure 2C). On the other hand, both the DcEntA- and DcEntB-expressing worms were also significantly more susceptible to infection by *D. coniospora* than control worms (Figure 2D). Thus, expression of certain individual fungal enterotoxins in the epidermis, albeit at an elevated level (Supplementary Figure S3), is sufficient to reduce *C. elegans* longevity and resistance to infection.

**Figure 2.**
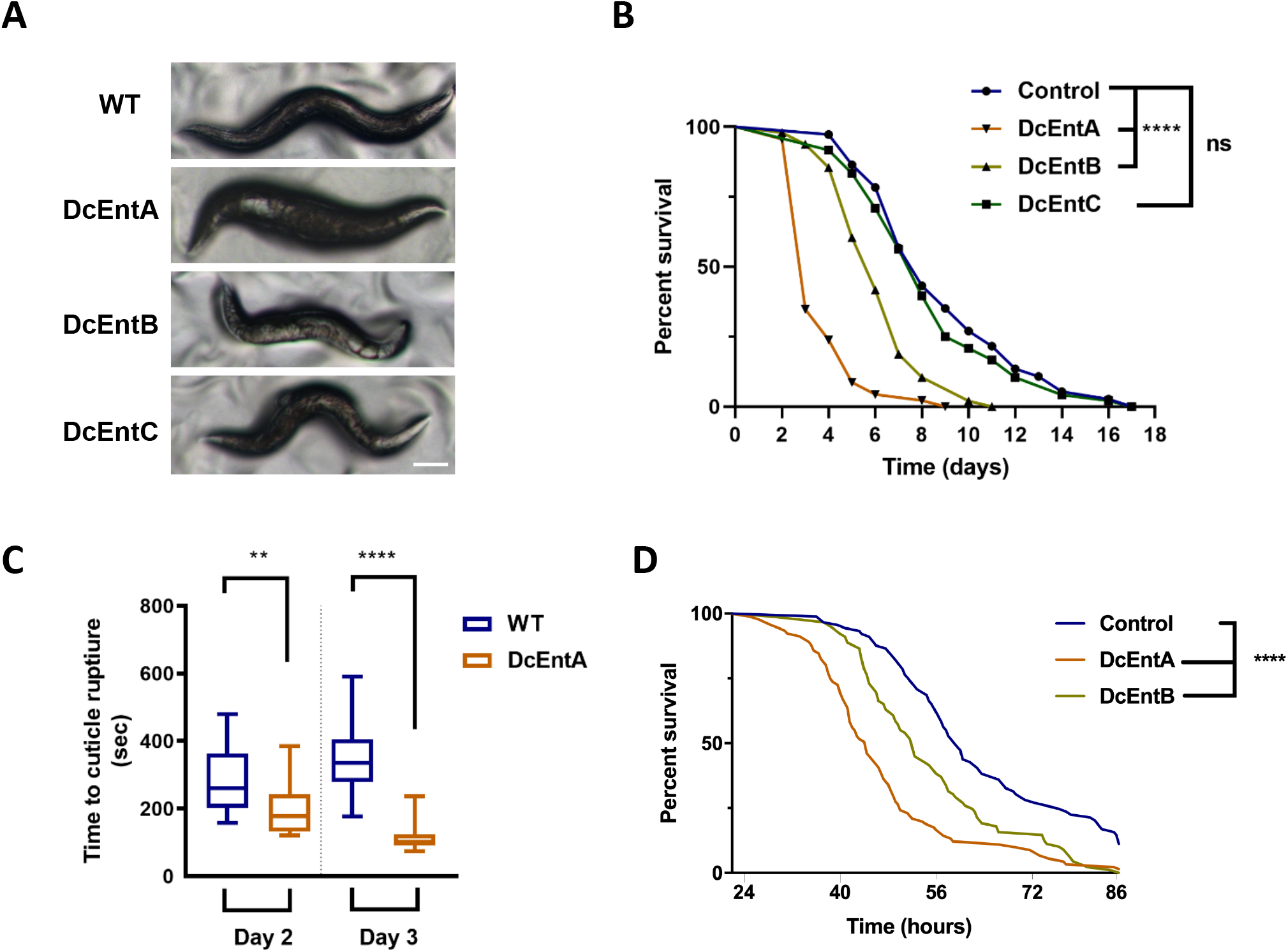
Expression of DcEntA or DcEntB makes worms sick and die precociously. (**A**) Representative images of control and enterotoxin-expressing worms on the third day of adulthood. From the top: control, DcEntA-, DcEntB- and DcEntC-expressing worms (JDW141, IG1926, IG1925 and IG1880, respectively); scale bar, 100 μm. (**B**) Lifespan counted from the L4 stage at 25°C of worms of these 4 strains. For each strain, n=50. **** p< 0.0001, one-sided log rank test. (**C**) Expression of DcEntA increases cuticle fragility in 2- and 3-day old adult worms. Tukey boxplots (n > 20, for each condition); unpaired t test, ** p < 0.01; **** p < 0.0001. (**D**) Survival of worms carrying *frIs7* and *hygR* transgenes (control; IG1864) or also expressing DcEntA (IG1942) or DcEntB (IG1941) after infection as young adults with *D. coniospora* at 25°C (n = 91, 89 and 89 respectively). Representative of 3 independent biological replicates.

### DcEntA blocks AMP gene expression after infection, downstream of a canonical p38 MAPK pathway

One of the key elements in the innate immune response of *C. elegans* to *D. coniospora* is the production of a battery of antimicrobial peptide (AMP) genes, including the well-studied *nlp-29* (Couillault et al., 2004; Pujol et al., 2012; Pujol et al., 2008b). When we assayed the effect of DcEntA expression on the induction of an *nlp-29p::GFP* reporter that is normally provoked by *D. coniospora* infection, we observed an almost complete block, that was not seen in worms expressing DcEntC (Figure 3A, B). Using qRT-PCR, we confirmed the inhibitory effect of DcEntA on *nlp-29* expression after infection, as well as on the expression of several other AMP genes of the *nlp* and *cnc* families (Figure 3C). Thus DcEntA blocks AMP gene expression at the transcriptional level.

**Figure 3.**
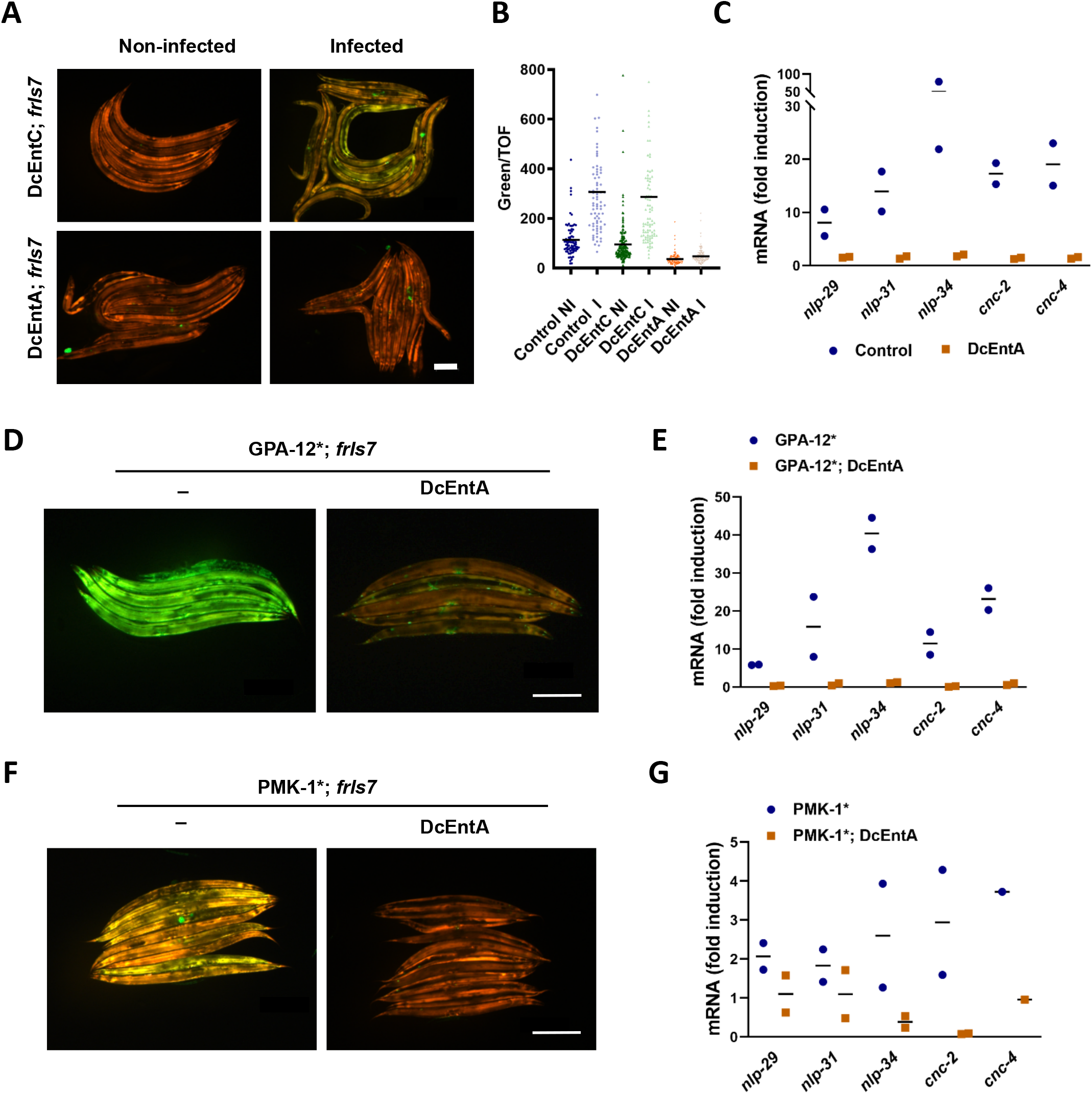
DcEntA blocks AMP gene expression after infection. (**A**) Representative images of DcEntC;*frIs7* (IG1883; upper panels) and DcEntA;*frIs7* (IG1942; lower panels) worms either not infected (left), or 20 h after infection with *D. coniospora* (right) as young adults. Red and green florescence is visualized simultaneously. Scale bar, 200 μm. (**B**) Quantification of relative green fluorescence in worms carrying *frIs7* and *hygR* transgenes (control; IG1864) or also expressing DcEntA (IG1942) or DcEntC (IG1883) either not infected (NI) or 20 h after infection (I) as young adults with *D. coniospora* at 25°C. (**C**) Quantitative RT-PCR analysis of the expression of *nlp* and *cnc* genes in worms carrying *frIs7* and *hygR* transgenes (control; IG1864) or also expressing DcEntA (IG1942) after 18h of infection by *D. coniospora*. Results from 2 independent experiments are shown relative to the expression levels in age-matched uninfected worms. (**D-G**) DcEntA acts downstream of *gpa-12* and *pmk-1* to block AMP gene expression. (**D**, **F**) Representative images of young adult worms carrying *frIs7*, expressing a constitutively active Gα protein (GPA-12*, **D**) or uncleavable p38 MAPK (PMK-1*, **F**) and expressing (right, IG1948 and IG1963) or not (left, IG1389 and BPW24) DcEntA. Red and green florescence is visualized simultaneously. Scale bar, 200 μm. (**E**, **G**) Quantitative RT-PCR analysis of the expression of *nlp* and *cnc* genes in worms expressing a constitutively active Gα protein (GPA-12* (IG1389), **E**) or uncleavable p38 MAPK (PMK-1* (BPW24), **G**) and worms also expressing DcEntA (DcEntA;GPA-12* (IG1948), **E**; DcEntA;PMK-1* (IG1963), **G**). Results from 2 independent experiments are shown relative to the expression levels in age-matched IG1864 worms.

The main pathway regulating *nlp-29* expression upon infection has been delineated (Kim et al., 2016; Polanowska et al., 2018; Zugasti et al., 2016). It starts with activation of the G-protein coupled receptor DCAR-1 by the endogenous ligand HPLA (Zugasti et al., 2014) and signal transduction via the Gα protein GPA-12 (Ziegler et al., 2009). Expression of a constitutively active form of the latter (referred to as GPA-12*) recapitulates many of the transcriptional changes that accompany infection (Lee et al., 2018), including an increased expression of *nlp-29*, and also leads to a high level of expression of the *nlp-29p::GFP* reporter (Labed et al., 2012). When we crossed the *DcEntA* transgene into a strain expressing GPA-12*, we observed an abrogation of the high *nlp-29p::GFP* reporter expression, and of the elevated expression of endogenous AMP genes as judged by qRT-PCR (Figure 3D, E). This indicates that DcEntA acts downstream of *gpa-12* to block defence gene expression.

The *gpa-12* pathway feeds into a conserved p38 MAPK cascade that ends with *pmk-1* (Pujol et al., 2008a). The activity of p38 MAPK PMK-1 is normally limited by proteolysis by the caspase CED-3. Mutating the caspase cleavage site (changing Asp327 to Glu) in PMK-1 results in higher p38 activity and increased expression of *nlp-29p::GFP* (Weaver et al., 2020). When we crossed the *DcEntA* transgene into a strain carrying the *pmk-1* (*D327E*) gain-of-function allele, we also observed a block of expression of the elevated expression of *nlp-29p::GFP* reporter and of endogenous AMP genes, although the effect was less dramatic than for the strain expressing GPA-12* since, as previously reported (Weaver et al., 2020), constitutive levels of *nlp* and *cnc* gene expression were only moderately elevated in the *pmk-1*(*D327E*) strain (Figure 3F, G). This indicates that DcEntA acts downstream of *pmk-1* to block defence gene expression.

### DcEntA affects the key immune regulators SNF-12 and STA-2

The p38 MAPK PMK-1 acts upstream of STA-2, a STAT-like protein. STA-2 is the common transcriptional regulator of *nlp* and *cnc* AMP genes. It interacts physically and functionally with the SLC6 protein SNF-12 (Dierking et al., 2011). When we crossed the *DcEntA* transgene into a strain expressing a SNF-12::GFP reporter protein, we observed a progressive disruption of its normal vesicular pattern at the apical surface of the hyp7 epidermal syncytium (Figure 4A, B). Since SNF-12 is essential for the induction of AMP gene expression, this disruption could be the cause of DcEntA’s inhibitory effect. Expression of DcEntA also significantly reduced the amount of a STA-2::GFP reporter protein within the nucleus (Figure 4C-E). Thus DcEntA could abrogate *nlp* and *cnc* AMP gene expression by preventing the accumulation of STA-2 in the nucleus upon infection, potentially indirectly, through an effect on membrane trafficking.

**Figure 4.**
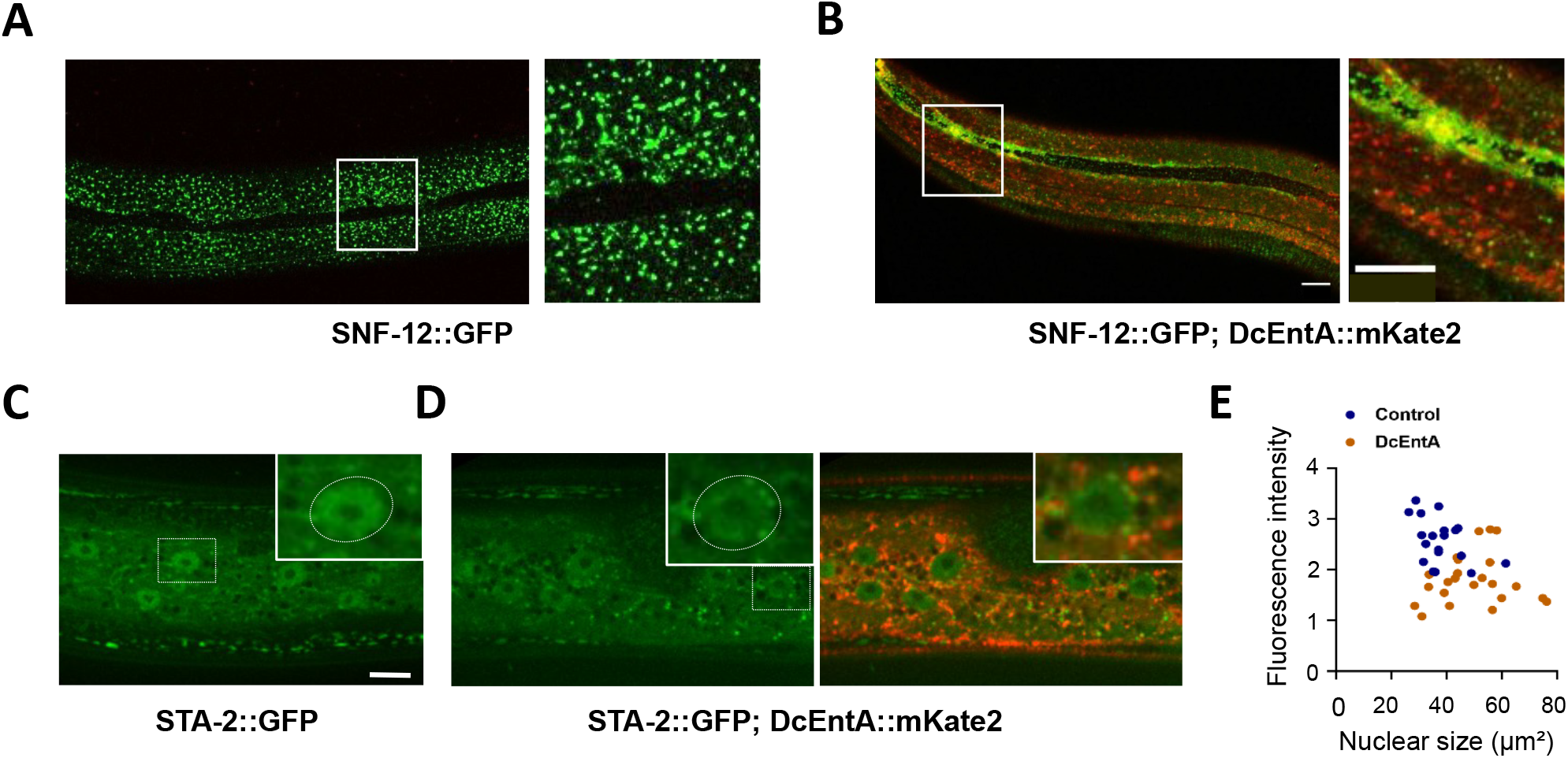
DcEntA alters the localisation of key immune regulatory proteins. (**A**, **B**) DcEntA disrupts the vesicular pattern of SNF-12. Confocal images, with a magnified view of the boxed area, of young adult worms expressing SNF-12::GFP alone (IG823, **A**) or together with DcEntA::mKate2 (IG1998, **B**) in the epidermis. Scale bar, 10 μm. (**C-E**) DcEntA decreases STA-2 nuclear accumulation. Confocal images of young adult worms expressing STA-2::GFP alone (XW18234, **C**) or with DcEntA::mKate2 (IG1971, **D**; left panel green channel only, right panel green and red channels shown together). STA-2::GFP levels are reduced in the epidermal nucleus (white box, enlarged insert; highlighted by white oval) in the presence of DcEntA. Scale bar, 20 μm. (**E**) The fluorescence intensity of STA-2::GFP in the nuclei of control (XW18234, blue) and DcEntA; STA-2::GFP worms (IG1971, brown) plotted against nuclear size, measured using ImageJ. Each dot represents a nucleus; n > 20. The difference between the 2 populations is significantly different (p < 0.0001; unpaired t-test).

### DcEntA can also provoke defence gene expression

Some genes, including *F40H7.12*, are induced upon *D. coniospora* infection but are not regulated by *gpa-12* (Lee et al., 2018). This nematode-specific gene, which encodes a protein of unknown function, shares many of the characteristics of “Intracellular Pathogen Response” genes (Reddy et al., 2017), including regulation by *pals-22* and *pals-25* (Reddy et al., 2019), and the fact that like *irg-1* and *irg-2* (Dunbar et al., 2012), its expression is strongly induced by the translational elongation inhibitor cycloheximide (CHX; Figure 5A). Thus upon *D. coniospora* infection the expression of *F40H7.12* is potentially increased in response to the consequences of infection, specifically a block in translation. This capacity to detect perturbation of normal protein synthesis has been reported to be an important part of the response of *C. elegans* to *Pseudomonas aeruginosa* infection (Dunbar et al., 2012; McEwan et al., 2012). It has thus far been unexplored in the context of *D. coniospora* infection; the expression of neither *irg-1* nor *irg-2* is induced after 12 or 24 hours of infection by *D. coniospora* (Engelmann et al., 2011). Notably, the expression of *F40H7.12*, as well as of *irg-1*, was markedly increased by DcEntA (Figure 5B, Supplementary Figure S4A). In contrast, *hsp-4*, a gene upregulated as part of the endoplasmic reticulum unfolded protein response (UPR), *hsp-6* and *hsp-60* that are markers of the mitochondrial UPR, as well as *gst-4* (oxidative stress) and *gpdh-1* (osmotic stress) (Calfon et al., 2002; Jones et al., 1989; Lamitina et al., 2006; Link and Johnson, 2002; Yoneda et al., 2004), showed no increase of expression in worms expressing DcEntA (Supplementary Figure S4B). Together these results indicate that DcEntA exerts a specific inhibitory effect on the expression of defence genes that are regulated by *snf-12* and *sta-2*, and far from acting via a generalized suppression of gene expression, actually triggers at least one infection-regulated gene, possibly because of an action blocking translation.

**Figure 5.**
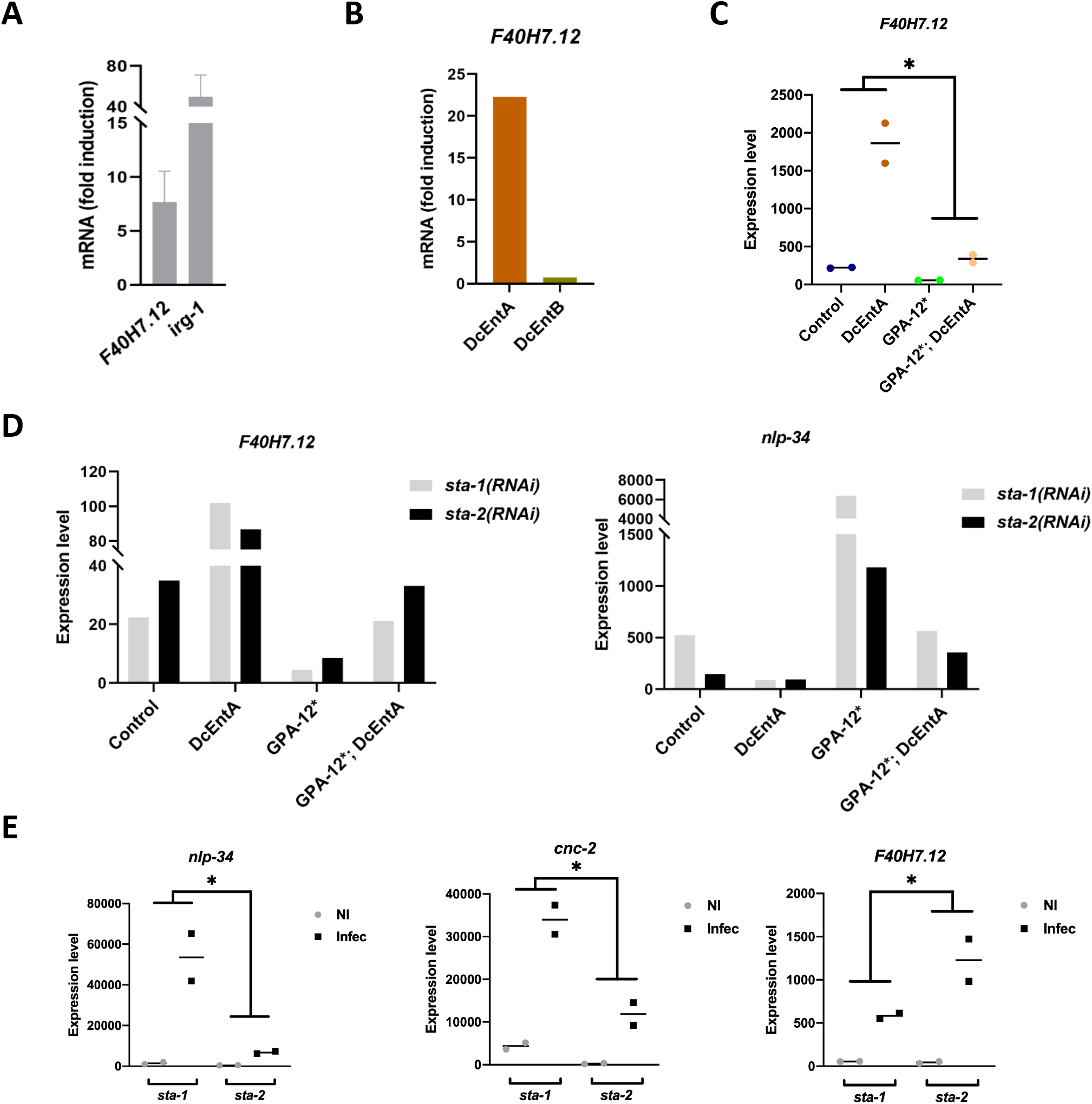
DcEntA induces expression of *F40H7.12*, a target of negative regulation by *sta-2*. (**A**) Quantitative RT-PCR analysis of the expression of *F40H7.12* and *irg-1* genes in worms carrying *frIs7* (IG274) following exposure as young adults to CHX for 6 h. Results from 3 independent experiments are shown as averages with standard deviation, relative to the expression levels in age-matched control worms. (**B**) Quantitative RT-PCR analysis of the expression of *F40H7.12* in young adult worms expressing DcEntA (IG1926) or DcEntB (IG1925). Results are presented relative to control worms (JDW141). (**C**) Quantitative RT-PCR analysis of the expression of *F40H7.12* in control (*hygR;frIs7* IG1864) worms and worms carrying *frIs7* expressing DcEntA (IG1942), GPA-12* (IG1389), or GPA-12*;DcEntA (IG1948). Data from two independent experiments are shown. The decrease in *F40H7.12* expression in IG1389 compared to IG1864 is consistent with previous results (Lee et al., 2018). (**D**) Quantitative RT-PCR analysis of the expression of *F40H7.12* and *nlp-34* in control (*hygR;frIs7* IG1864) worms and worms expressing DcEntA (IG1942), GPA-12* (IG1389), and GPA-12*;DcEntA (IG1948) following RNAi against *sta-1* or *sta-2*. See Supplementary Figure S5 for two further replicates. (**E**) Quantitative RT-PCR analysis of the expression of *nlp-34*, *cnc-2* and *F40H7.12* following RNAi against *sta-1* or *sta-2* in worms infected for 18h and non-infected (NI) controls in the epidermis-specific RNAi strain IG1502. Data from two independent experiments are shown. Here and in (C), the fold-change in expression level between the 2 indicated conditions is significantly different (* p < 0.05; paired one-sided t test).

### DcEntA further promotes defence gene expression by removing a STA-2-dependent repression

To investigate whether there was any cross-talk between the pathway regulating *F40H7.12* and the well-studied p38 MAPK pathway regulating *nlp-29* expression, we assayed *F40H7.12* expression in the strain expressing GPA-12* and DcEntA. There was a marked decrease in its expression, compared to the DcEntA strain (Figure 5C). In vertebrates, as well as acting as positive regulators of gene expression, STAT proteins can directly repress the accessibility and transcription of specific loci (Mandal et al., 2011). Indeed, the only other STAT protein in *C. elegans*, STA-1, is known to be a repressor of virus infection response genes (Tanguy et al., 2017). In a parsimonious model, the effect of GPA-12* could be mediated by STA-2, acting as a negative regulator of *F40H7.12*. Consistent with this, in control animals, and in animals expressing GPA-12* alone or in combination with DcEntA, *sta-2(RNAi)* indeed led to an increase in *F40H7.12* expression. On the other hand, *sta-2(RNAi)* did not increase *F40H7.12* expression in worms expressing DcEntA alone (Figure 5D, Supplementary Figure S5). These results can be explained if one assumes that when the p38 MAPK pathway is not active, DcEntA almost completely inhibits STA-2 function, so *sta-2(RNAi)* cannot have an effect, but this is partially countered in the GPA-12* background. Such an interpretation of opposing regulatory effects was supported by an analysis of the expression of the *sta-2*-dependent AMP gene *nlp-34*, wherein the pattern of changes in its level was the inverse of that seen for *F40H7.12* (Figure 5D, Supplementary Figure S5). In further support, when we knocked down *sta-2* specifically in the epidermis (in strain IG1502), and then infected worms with *D. coniospora*, we observed a significant increase in the induction of *F40H7.12*, and the expected decrease for the AMP genes *nlp-34* and *cnc-2* (Figure 5E). Thus, by preventing accumulation of STA-2 in the nucleus, DcEntA blocks both its transcriptional activator and repressor functions.

### DcEntA has ADP-ribosylation activity and potentially interacts with many host proteins

As a first step to understanding the basis of these effects, we took an unbiased biochemical approach to identify proteins that interacted with DcEntA *in vivo*. From a synchronized population of adult worms expressing the DcEntA fusion protein, we pulled down DcEntA::FLAG::Degron::mKate2 by immunoprecipitation from whole worm extracts and subjected the purified proteins to mass spectrometry analysis. Samples from four other strains of transgenic worms, each expressing a different candidate virulence factor, were processed in parallel (see below; Harding *et al*., in preparation) allowing proteins that specifically interacted with DcEntA to be identified. As expected, DcEntA itself featured among the most abundant proteins identified (Figure 6, Supplementary Table S5). Detailed examination of its spectrum supported the presence of at least one site with an ADP-ribosylation modification, at an asparagine in the non-conserved C-terminus of the protein (Supplementary Figure S6). This suggests that in common with some bacterial exotoxins (Krueger and Barbieri, 1995), DcEntA is able to auto-ADP-ribosylate.

**Figure 6.**
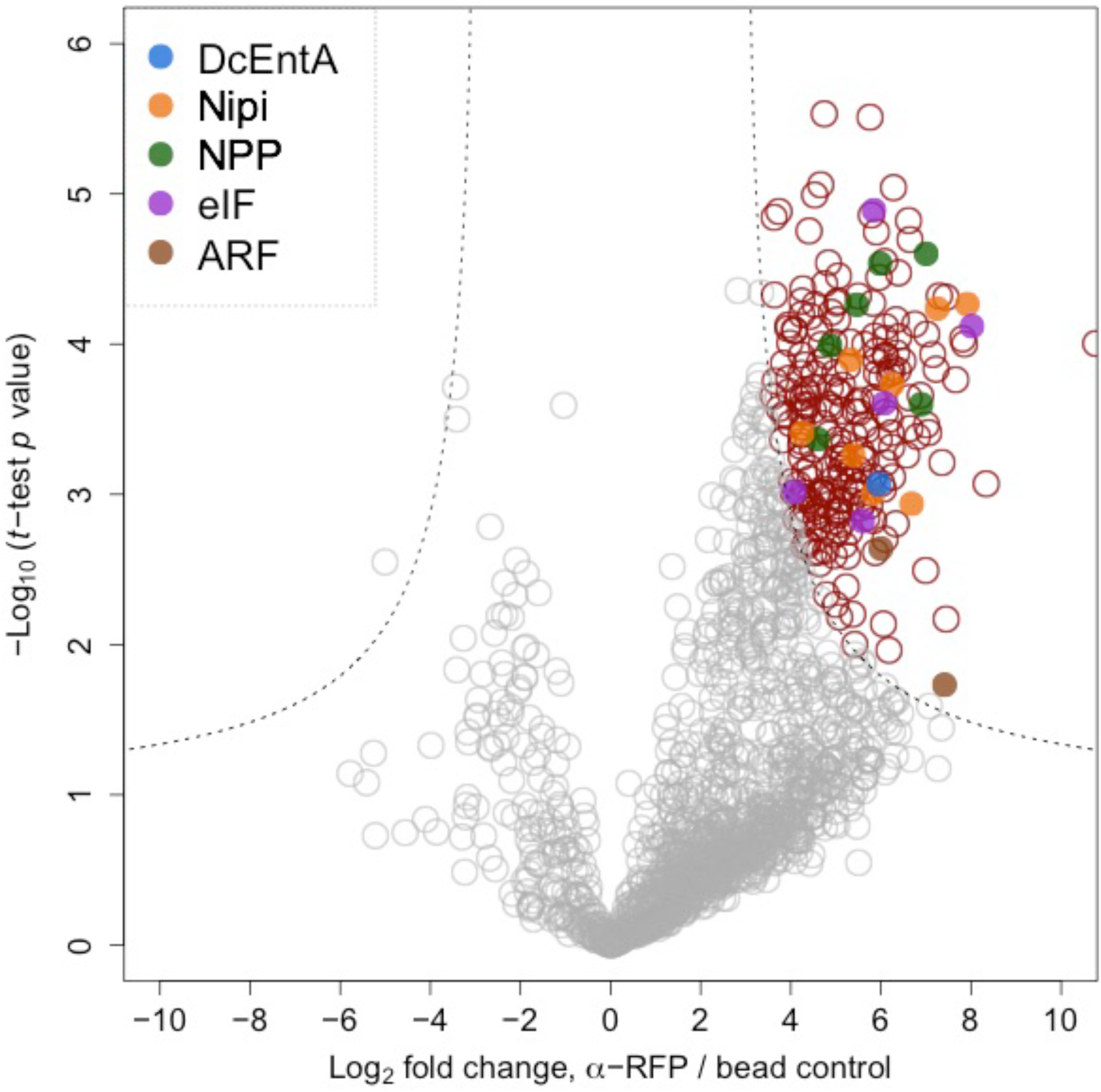
DcEntA interactors identified by label-free quantitative immunoprecipitation. The relative abundance of proteins co-precipitated with DcEntA::FLAG::Degron::mKate2 was assessed by mass spectrometry. Volcano plot showing specific interaction partners (in red) of DcEntA::FLAG::Degron::mKate2 (DcEntA in blue). The mean values for fold change from 3 independent experiments are shown. The SAM (significance analysis of microarrays) algorithm was used to evaluate the enrichment of the detected proteins. Proteins that met the combined enrichment threshold (hyperbolic curves, *t*_0_ = 2) are coloured in red. The 2 *C. elegans* ARF proteins are shown in brown, known members of the nuclear pore complex (NPP) are in green, eukaryotic initiation factor proteins (eIF) in purple and proteins corresponding to Nipi (No Induction of Peptide after *Drechmeria* Infection) genes in orange.

Neither SNF-12 nor STA-2 were found among the specific interactors of DcEntA, suggesting that its effects on their localisation are likely to be indirect (see below). Among the 245 candidates identified through mass spectrometry, 2 proteins stood out. Cholera toxin from *Vibrio cholera* requires a host protein, ADP ribosylation factor (ARF), in order to ADP-ribosylate its targets (Kahn and Gilman, 1986; Moss and Vaughan, 1995). The 2 nematode orthologues of the human ARF1 and ARF3, ARF-1.1 and ARF-1.2, were found among the interactors of DcEntA, consistent with a conserved mode of action for this fungal heat-labile enterotoxin.

### DcEntA potentially affects diverse aspects of host cell physiology

To get an overall view of the candidate proteins, we used the gene set enrichment analysis tools available within Wormbase (Angeles-Albores et al., 2018). Among the enriched phenotype classes, given the disruptive effect of DcEntA on the vesicular pattern of SNF-12, “vesicle organization variant” (WBPhenotype:0001671; p=5.4×10^−3^) stood out. The most highly over-represented gene ontology term was the “Cellular Component” class “ribonucleoprotein granule” (GO:0035770; p=5.6 ×10^−6^). This class is for components of a “non-membranous macromolecular complex containing proteins and translationally silenced mRNAs”. This is consistent with the observed increase in *irg-1* expression, a marker of translation inhibition (Dunbar et al., 2012; McEwan et al., 2012), provoked by DcEntA, and the enrichment for “peptide biosynthetic process” (GO:0043043; p=1.2×10^−4^) and several other translation-related classes (Supplementary Table S5). DcEntA also disrupts membrane trafficking and we found an enrichment in the classes “cellular macromolecule localization” (GO:0070727; p=1×10^−4^) and “nuclear outer membrane-endoplasmic reticulum membrane network” (GO:0042175; p=1.6×10^−4^). Both classes included COGC-3, part of the Conserved Oligomeric Golgi (COG) Component, a peripheral membrane 8-protein complex that acts in intra-Golgi trafficking. COGC-2 and COGC-8 were also found as specific interactors (but are not annotated with GO:0042175 or GO:0070727), suggesting that DcEntA might interfere with Golgi trafficking. Several other components of *C. elegans* intracellular vesicle transport machinery were identified, including 16 that have been reported to interact physically with VPS-45, orthologue of human VPS45 (vacuolar protein sorting 45 homolog), required for RAB-5-dependent endocytic transport (Gengyo-Ando et al., 2007), and 11 interactors of LET-413 (Waaijers et al., 2016), the nematode Erbin protein that acts as a RAB-5 effector during endocytic recycling (Liu et al., 2018). The list of candidate DcEntA interactors also included several proteins involved in cytoskeleton dynamics, including the nematode ezrin/moesin/radixin orthologue ERM-1, the microtubule plus-end binding protein EBP-1, and the twinfilin homologue TWF-2, an actin binding protein. Coordinated changes in microtubule and actin dynamics are required for the proper recruitment of SNF-12 to sites of injury in hyp7. Without SNF-12 recruitment, there is an abrogation of the immune response (Taffoni et al., 2020). There are thus several potential ways that DcEntA could influence membrane trafficking and SNF-12 localisation. Interestingly, given the pattern of SNF-12 observed upon expression of DcEntA, with an enrichment along the baso-lateral seam cell boundary, 38 of its interactors are known to be potential binding partners of DLG-1, which is found in the same location (Waaijers et al., 2016). We also identified 11 proteins reported to interact with AKIR-1, which is essential for AMP gene expression in hyp7 (Polanowska et al., 2018), suggesting an additional way in which DcEntA might affect the host response. There were also 6 NPP proteins, components of the nuclear pore, required for nucleocytoplasmic transport, and at least 5 involved in translation initiation. Two of each category (NPP-1 and NPP-6; EIF-2gamma and F33D11.10, respectively) had previously been implicated in the regulation of AMP gene expression: the 4 corresponding genes were identified in a whole-genome RNAi screen for Nipi genes, positive regulators of *nlp-29* expression (Zugasti et al., 2016). Another 8 candidate DcEntA interactors correspond to Nipi genes (Figure 6, Supplementary Table S5). Thus it is possible that DcEntA has multiple modes of action, altering membrane trafficking and thus the function of SNF-12, potentially blocking nuclear import of STA-2 via alteration of the nuclear pore thereby preventing AMP gene transcription, as well as affecting translation of host defence genes. Taken together, these results lead to a model in which the expression of DcEntA, acting in concert with the host Arf GTPases, through ADP ribosylation, directly or indirectly prevents accumulation of STA-2 in the nucleus and thereby inhibits the expression of multiple defence genes, rendering *C. elegans* more susceptible to infection. At the same time, potentially as a counter-defensive mechanism, loss of STA-2-dependent repression accentuates the increase in the expression of *F40H7.12* provoked by *D. coniospora*, and presumably of other host genes that potentially help protect against infection and that are induced following a reduction in translation (Figure 7).

**Figure 7.**
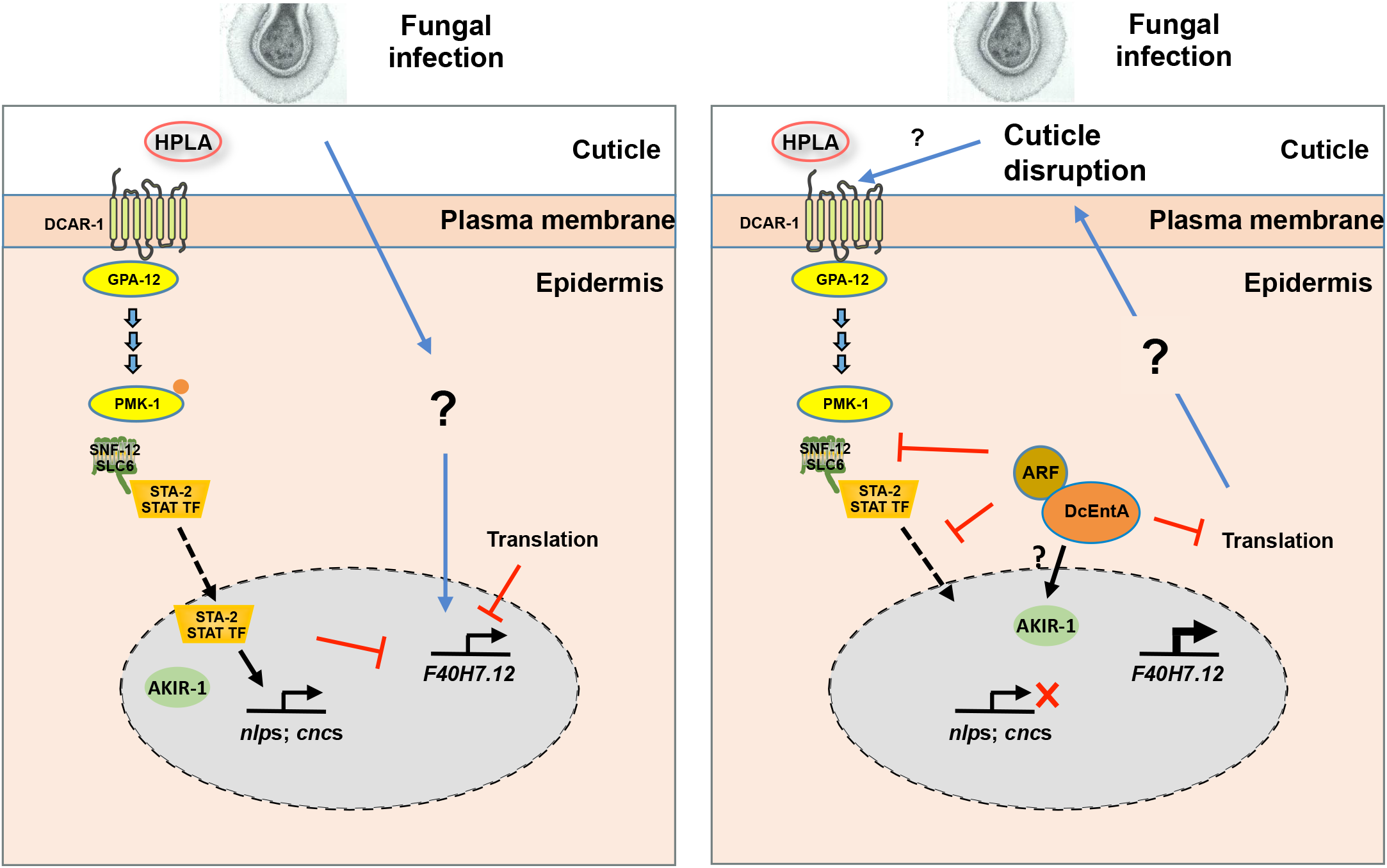
Models of the innate immune response to *D. coniospora* infection and DcEntA action. Left panel: Infection by *D. coniospora* causes an increase in HPLA, activation of its receptor DCAR-1 and, via GPA-12, the p38 MAPK (PMK-1) cascade. PMK-1 acts upstream of SNF-12/STA-2. The subsequent translocation of STA-2 into the nucleus leads to an increase in the expression of *nlp* and *cnc* genes, which depends also on the AKIR-1 complex. An uncharacterized parallel pathway, potentially linked to surveillance of translation, activates *F40H7.12*, which is also negatively regulated by DCAR-1/STA-2 pathway. Right panel: DcEntA, via its interaction with ARF proteins and other protein partners (brown circle), potentially interferes with the normal innate immune response at multiple levels. It alters SNF-12 localization, blocks STA-2 nuclear translocation and AMP gene expression, and possibly interacts with the AKIR-1 complex. It may inhibit translation, leading to an increase in *F40H7.12* expression, accentuated by the loss of the repressive function of STA-2. DcEntA causes increased cuticle fragility, possibly through inhibition of translation. This has the potential to increase HPLA levels, adding a further level of complexity to the system.

### DcEntB affects nucleolar size and shape

Turning to DcEntB, as described above, it localises to the nucleolus. Nucleoli are the site of ribosome biogenesis and also play a role in cells’ responses to diverse stresses (Latonen, 2019). They have been linked to the regulation of immune responses against bacterial pathogens in *C. elegans* (Fuhrman et al., 2009; Tiku et al., 2018). Interestingly, we observed a clear alteration of nucleolar morphology in worms expressing DcEntB, with many nucleoli exhibiting strikingly angular shapes rather than their usual spherical form (Figure 8A, B). This is reminiscent of the morphological changes that accompany several different pharmacological treatments, including ATP depletion, in mammalian cells (Caragine et al., 2019). Expression of DcEntB was associated with an increase in nuclear size (Figure 8C), and a proportionately larger increase in the size of nucleoli (Figure 8D). We also used the FIB-1::GFP reporter (Yi et al., 2015) to characterise and quantify these changes, and confirmed a highly significant increase in nucleolus size and Feret's diameter. The latter is a measure of maximum length and when compared to area gives an indication of any deviation from circularity (Figure 8E, F, Supplementary Figure S7A).

**Figure 8.**
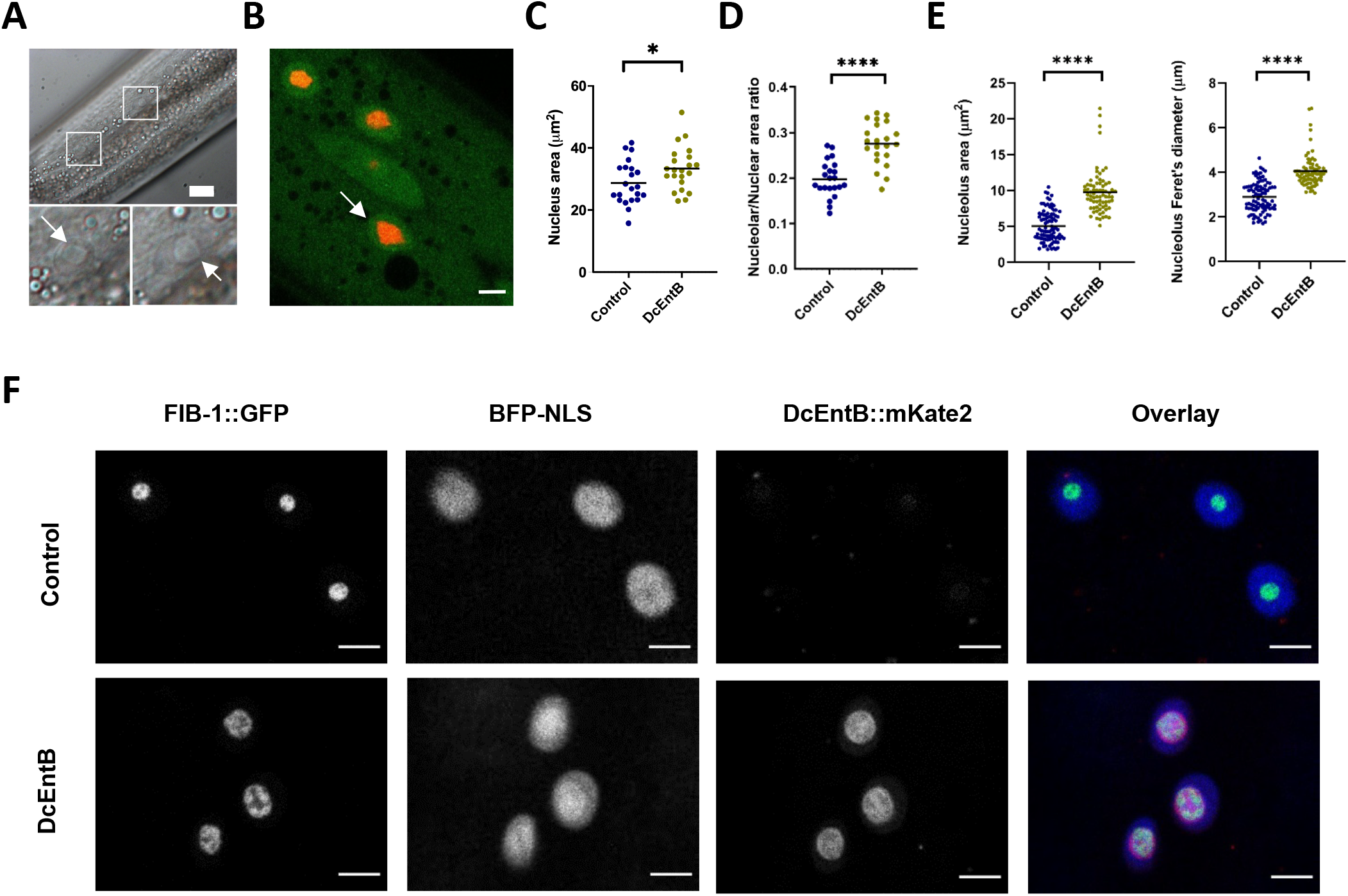
DcEntB makes nucleoli irregular and larger. (**A**) Representative Normarski image of a young adult worm expressing DcEntB. Scale bar, 10 μm. Enlarged views of the boxed regions show the large and irregular nucleoli (white arrows). (**B**) Representative confocal image of a young adult worm expressing DcEntB::mKate2 and STA-2::GFP (IG1977) in hyp7. White arrow points to an irregular nucleolus (red). Scale bar, 10 μm. (**C**, **D**) Quantification of nuclear size (**C**) and nucleolus/nucleus ratio (**D**) in the epidermis of young adult IG1977 worms (DcEntB) and their siblings without the DcEntB transgene (control). n > 20 in each strain; bars represent the mean; * p < 0.05, unpaired t-test **** p < 0.0001, unpaired t-test. (**E**) Quantification of nucleolus area and Feret’s diameter in hyp7 of young adult worms expressing FIB-1::GFP with (DcEntB, IG1984) or without (control, SJL1) DcEntB. **** p < 0.0001. Statistical significance was determined using a nonparametric Mann Whitney test. (**F**) Confocal images of hyp7 nuclei in young adult worms expressing FIB-1::GFP with (DcEntB, IG1984; lower panels) or without (control; SJL1; upper panels) DcEntB, scale bar, 5 μm.

### DcEntB increases AMP gene expression and can prime the immune system

In direct contrast to the effect of DcEntA, expression of DcEntB was associated with an increase in the constitutive expression of the *nlp-29p::GFP* reporter (Figure 9A). This increase was dependent upon STA-2 since *sta-2*(*RNAi*) reduced reporter gene expression back to the normal level (Figure 9B). Using qRT-PCR, we confirmed the positive effect of DcEntB on *nlp-29* expression, and demonstrated a similar *sta-2*-dependent effect for several *nlp* and *cnc* genes (Figure 9C). Notably, and confirming our previous result (Figure 5B), worms expressing DcEntB did not exhibit any change in *F40H7.12* expression (Figure 9C). Together, these results suggest that DcEntB might act through the canonical STA-2 pathway to regulate AMP gene expression. Supporting this, when we crossed the DcEntB transgene into a strain expressing STA-2::GFP, we observed a significant increase in the amount of STA-2::GFP in the nucleus (Figure 9D; p < 0.0001, unpaired t-test).

**Figure 9.**
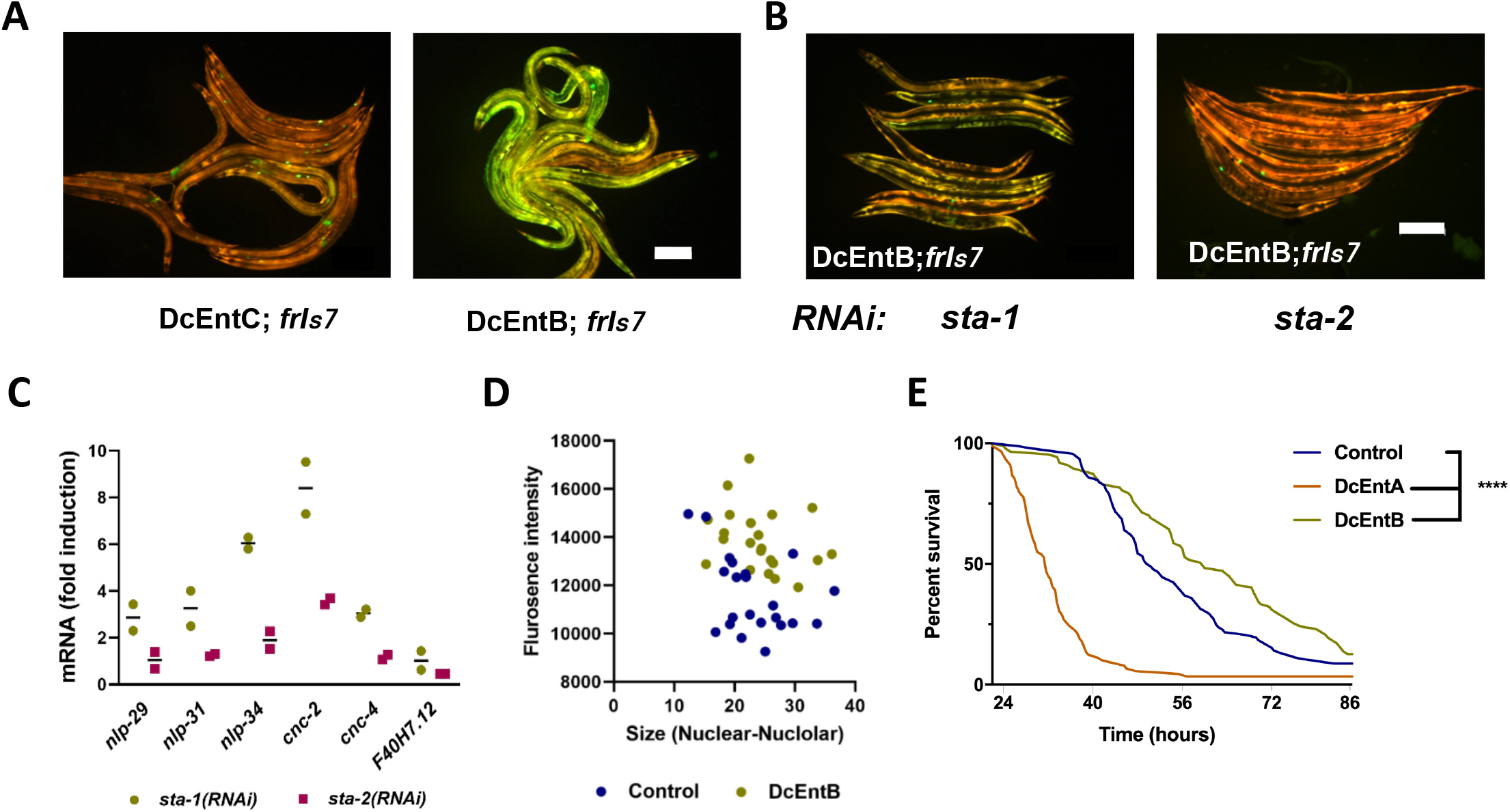
DcEntB induces AMP gene expression in a *sta-2*-dependent manner. (**A**, **B**) Representative images of adult worms, 3 days after the L4 stage, carrying *frIs7* and expressing DcEntC (IG1883) or DcEntB (IG1941) on *E. coli* OP50 (**A**) or following RNAi against *sta-1* or *sta-2* (**B**). Red and green florescence is visualized simultaneously. Scale bar, 200 μm. A difference in GFP levels for worms with *frIs7* between OP50 and RNAi (HT115) bacteria has been observed regardless of the genetic background. (**C**) Quantitative RT-PCR analysis of the expression of *nlp*, *cnc* and *F40H7.12* genes in worms expressing DcEntB (IG1941) following RNAi against *sta-1* or *sta-2*. Results from 2 independent experiments are shown relative to the expression levels in age-matched control (IG1864) worms. (**D**) The fluorescence intensity of STA-2::GFP in the nuclei of control (XW18234, blue) and DcEntB; STA-2::GFP worms (IG1977, green) plotted against the nuclear-nucleolar size, measured using ImageJ. Each dot represents a nucleus; n>20. (**E**) Survival of control (IG1864) worms and worms expressing DcEntA (IG1942) or DcEntB (IG1941) after infection as young adults with a concentration of *D. coniospora* spores 10 times higher than usual at 25°C (n = 92, 91 and 87 respectively). The curves here are representative of 3 independent biological replicates.

There is therefore a striking dichotomy between the effects of DcEntA and DcEntB. DcEntA appears to act as many known virulence factors do, blocking the activation of an immune defence pathway, in this case potentially via an inhibition of the activity of the transcription factor STA-2. DcEntB, on the other hand, appears to activate the same pathway, leading to more STA-2 in the nucleus and more AMP gene expression. We hypothesised that this might reflect a host defence strategy wherein the presence of DcEntB is detected, either directly or indirectly, as a form of surveillance immunity. Should this be the case, one would predict that expression of DcEntB might increase survival following infection. Although we showed that following infection under standard conditions DcEntB-expressing worms had an increased susceptibility to infection (Figure 2D), when we infected the same strain of worms with *D. coniospora* using a very high concentration of spores, in contrast to worms expressing DcEntA that died more rapidly, the DcEntB-expressing worms were indeed significantly resistant, and lived longer even than the controls (Figure 9E). Thus the presence of high levels of DcEntB does appear to have the capacity to prime the host immune system.

### DcEntB-induced changes in nucleolar morphology require STA-2 and are associated with a specific induction of targets of the p38 MAPK pathway

To investigate the link between the observed changes in STA-2 nuclear occupancy, AMP gene expression and nucleolar morphology, we first assayed whether *sta-2*(*RNAi*) affected the shape of nucleoli, using the FIB-1::GFP reporter strain. In contrast to control worms, in worms expressing DcEntB, we observed a very marked decrease in the DcEntB-associated phenotypes, both size and Feret’s diameter, upon *sta-2*(*RNAi*) (Figure 10A, Supplementary Figure S7B), suggesting that the change in nucleolar morphology could be a consequence of DcEntB’s recruitment of STA-2 to the nucleus and/or the resulting STA-2-dependent changes in gene expression. Notably, when we quantified nucleolar morphology in the FIB-1::GFP reporter strain infected with *D. coniospora*, we observed a modest increase in size relative to uninfected controls, without a significant change in Feret’s diameter (Figure 10B, Supplementary Figure S8A).

**Figure 10.**
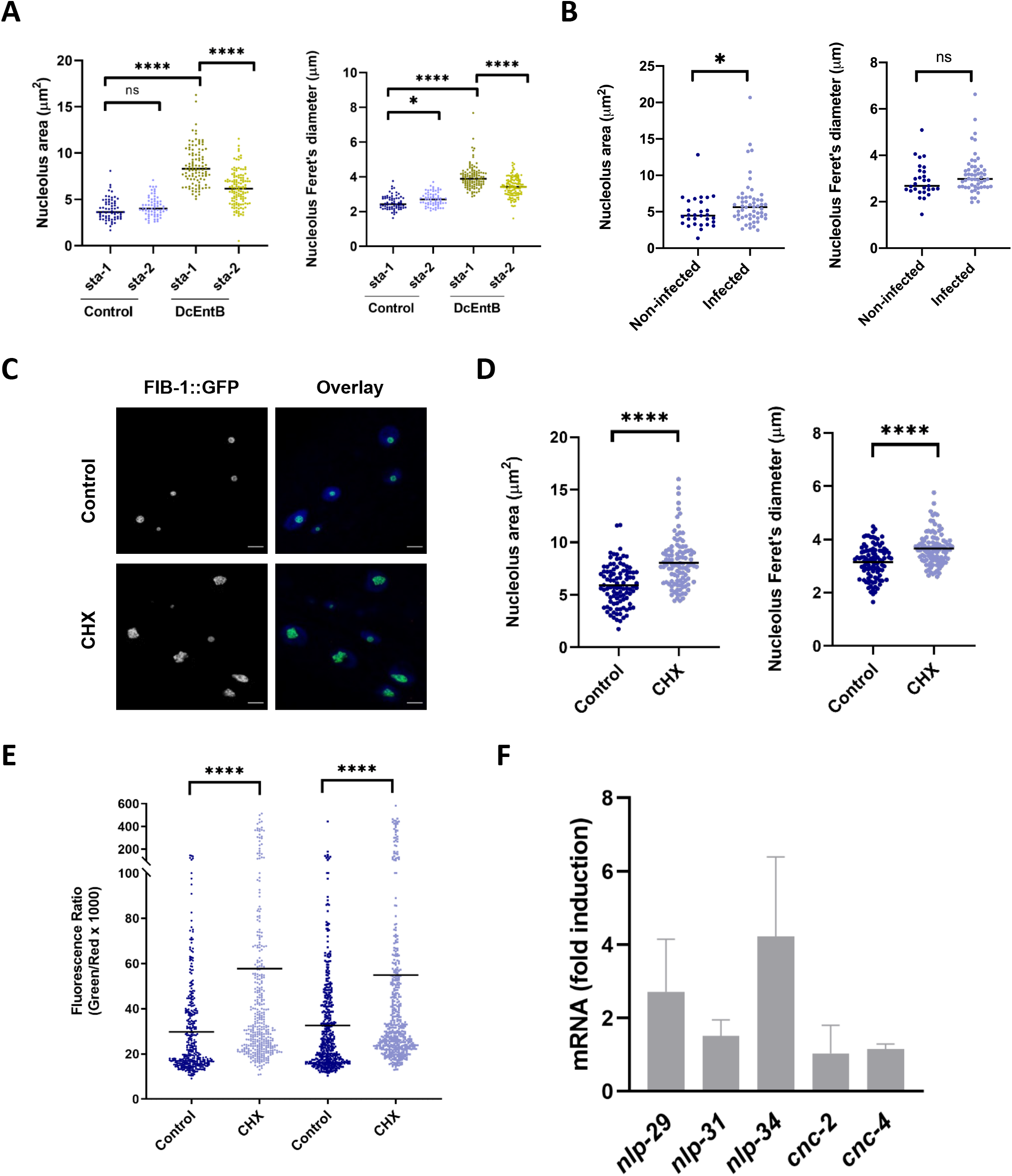
Relationship between nucleolar morphology, translation and AMP gene expression. (**A**, **B**) Quantification of nucleolus area and Feret’s diameter in young adult worms expressing FIB-1::GFP with (DcEntB; IG1984) or without (Control; SJL1) DcEntB following RNAi against *sta-1* or *sta-2* (**A**) or in SJL1 worms following 24 h infection (**B**). (**C**) Representative confocal images of young adult worms expressing FIB-1::GFP (SJL1) after exposure for 6 h to cycloheximide (CHX; lower panels) compared to control (upper panels), scale bar, 5 μm. (**D**) Quantification of nucleolus area and Feret’s diameter in young adult worms expressing FIB-1::GFP (SJL1) after exposure for 6h to CHX, compared to control. (**E**) Quantification of relative green fluorescence of young adult worms carrying *frIs7* (IG274) after exposure for 6 h to CHX, compared to control. The results from 2 independent experiments are shown. (**F**) Quantitative RT-PCR analysis of the expression of *nlp* and *cnc* genes in worms carrying *frIs7* (IG274) following exposure as young adults to CHX for 6 h. Results from 3 independent experiments are shown as averages with standard deviation, relative to the expression levels in age-matched control worms. Statistical significance was determined using a nonparametric Mann Whitney test; * p < 0.05, **** p < 0.0001, ns, not significant.

To explore further the relationship between nucleolar morphology and the *sta-2*-dependent immune response, we treated worms carrying FIB-1::GFP with the protein synthesis inhibitor CHX, a drug that is also known to alter nucleolar shape (Caragine et al., 2019). While exposure to 500 μg/ml CHX for prolonged periods affects development and animal health (Eberhard et al., 2013), we found that adults tolerated this concentration well for short (6 h) periods. It did, however, cause nucleoli to become larger and less round (Figure 10C, D, Supplementary Figure S8B). Additionally, it led to a modest but significant increase in the expression of the *nlp-29p::GFP* reporter (Figure 10E). Using qRT-PCR, we confirmed this effect on *nlp-29* expression, and demonstrated a similar effect for *nlp-34* (Figure 10E, F), albeit to a lesser degree than *irg-1*, which responds strongly to a block of protein synthesis (Figure 5A; (Dunbar et al., 2012)). Notably, the expression of *cnc-2* and *cnc-4* was not affected by CHX treatment (Figure 10F). These 2 genes are not regulated by p38 MAPK PMK-1 upon *D. coniospora* infection (Zugasti and Ewbank, 2009). Taken together with the fact that DcEntB does not increase the expression of *F40H7.12* (Figure 5B, Figure 9C), but CHX treatment does (Figure 5A), these results suggest that the potential inhibition of protein synthesis by DcEntB leads to a specific induction of a p38 MAPK dependent immune response in the epidermis, in addition to its more direct effect on STA-2 activity (Figure 11).

**Figure 11.**
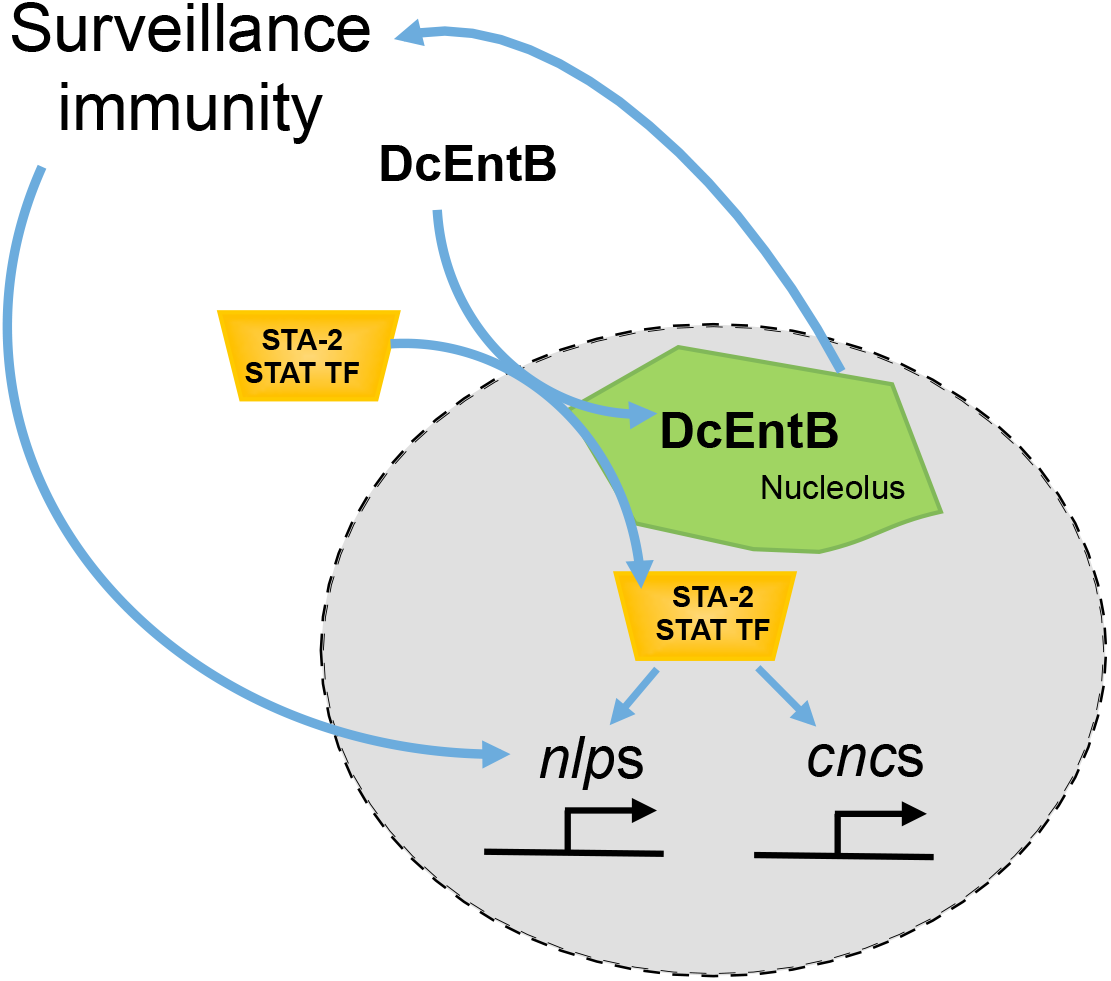
Model of DcEntB action. When DcEntB is expressed in hyp7, it localises to the nucleolus. Expression of DcEntB drives STA-2 into the nucleus, leading to an increase in *nlp* and *cnc* gene expression. DcEntB alters the shape and size of the epidermal nucleolus. This provokes a surveillance mechanism leading to the expression of *nlp* but not *cnc* genes.

### DcEntB potentially interacts with many host proteins to affect diverse aspects of host cell physiology

In an attempt to understand better the complex effects of DcEntB we undertook the same type of biochemical approach as described above for DcEntA, in the hope of finding meaningful protein partners. We identified 121 proteins as being specifically enriched in the proteins co-immunoprecipitated with DcEntB (Supplementary Table S6). Currently, just 150 *C. elegans* proteins have the Gene Ontology cellular component annotation “nucleolus” (GO:0005730). Only one of them, NST-1, orthologue of human GNL3L (G protein nucleolar 3 like) was among the candidate DcEntB interactors. GNL3L is a nucleolar GTPase that is essential for ribosomal pre-rRNA processing and cell proliferation. As mentioned above, many heat-labile enterotoxins exert their effects through ADP-ribosylation of guanine nucleotide-binding proteins. Whether NST-1 is an authentic interactor of DcEntB remains, however, to be established. When we used the NoD NoLS predictor (Scott et al., 2011), there were a further 17 potential nucleolar proteins (Supplementary Table S6). They included LET-502, homologue of Rho-associated coiled-coil kinase (ROCK).

Interestingly, it was recently shown that during human cytomegalovirus infection, ROCK1 translocates to the nucleus and concentrates in the nucleolus (Eliyahu et al., 2019), so this interaction merits investigation. The same applies to the other candidate interactors with GTP binding or GTPase activity, including PES-7/IQGAP, LET-60/HRas, RAB-6.1/RAB6A and CDC-42/CDC42, that are all predicted to have an NoLS (Supplementary Table S6).

Two of the top 6 potential protein partners have been identified as interacting with ATX-2/ATXN2L. ATX-2 associates directly with Rab GDP dissociation inhibitor, which is thought to regulate the activity of the Ras family protein RHEB and thereby the mTORC1 pathway (Bar et al., 2016). This in turn impacts the nucleolus in many different ways, including multiple steps of ribosome biogenesis (Iadevaia et al., 2014; Iadevaia et al., 2012; Raman et al., 2014). It should be noted, however, that while ATX-2 is found in both the nucleus and cytoplasm, and despite a NoD-predicted NoLS, it is absent from the nucleolus in *C. elegans* (Bar et al., 2016), so the relevance of these potential interactors is unclear.

One of the consequences of DcEntB expression is an increase in *nlp-29p::GFP* reporter gene expression. In a previous genome-wide RNAi screen for this same phenotype, we identified close to 300 genes (Zugasti et al., 2016). Among the corresponding proteins, only one, F10E7.5 is predicted to be nucleolar and it was not found in our current list. Only 2 proteins in the list correspond to candidate DcEntB interactors (SEM-5 and K08E3.5), but as neither is predicted to be nucleolar (Supplementary Table S6), again the relevance of these observations is unclear.

To take a broader view, we used the enrichment tools in Wormbase. In contrast to the DcEntA interactors, there were few enriched classes. Among them, “nuclei enlarged” (WBPhenotype:0001567; p=7.4×10^−4^) stood out, with both LET-502/ROCK and LET-60/HRas associated with this term ((Green et al., 2011); Supplementary Table S6). Given the alteration of nuclear size and nucleolar morphology provoked by DcEntB expression, these candidates merit further investigation. As knocking down the gene corresponding to one other candidate protein with this annotation, RNP-4, (orthologue of human RBM8A (RNA binding motif protein 8A) causes a block in AMP gene expression (Zugasti et al., 2016), understanding precisely the consequences of DcEntB expression will be challenging. It would be facilitated by identifying those host proteins that are not only interactors of DcEntB but also substrates for its enzymatic activity. For the time being, we did not find convincing evidence for specific ADP-ribosylation of any of the candidate enterotoxin interactors. ADP-ribose is a labile group that breaks easily with the fragmentation method used for the mass spectrometric analysis (Bonfiglio et al., 2017). Nevertheless, the results reported here represent an important step in understanding the complexity of the molecular interactions that underlie *D. coniospora*’s capacity to infect and kill *C. elegans*.

## DISCUSSION

The comprehension of fungal pathogenesis requires the identification of virulence factors and a dissection of their mode of action. In the current study, we chose to express individual candidate fungal virulence factors, as tagged chimeric proteins, at an elevated level, in a single tissue, the multi-nuclear epidermal syncytium hyp7. Tagging proteins can affect their function. For example, about 10% of the proteins in yeast are localized to different subcellular localizations when tagged at the carboxy terminus or amino terminus (Weill et al., 2019). There has, however, been sufficient experience with fluorescent reporter proteins, in *C. elegans* and in other model organisms, including genome-wide surveys of protein localization (Huh et al., 2003), to know that in most cases the chimeric protein acts like its unmodified counterpart. Our approach is analogous to a recent study where numerous candidate secreted effector proteins from the plant pathogen *Colletotrichum higginsianum* were expressed as N-terminal fusions with GFP directly inside plant cells and determined to localize to peroxisomes, Golgi bodies, and microtubules (Robin et al., 2018). As another example, a putative virulence factor from the nematode-trapping fungus *Duddingtonia flagrans* was expressed in *C. elegans* as a C-terminal GFP-fusion construct where it is localized to nuclei, consistent with the presence of an NLS in its sequence (Youssar et al., 2019). Among the *D. coniospora* proteins we studied here, DcEntB had a nucleolar localization, as predicted *in silico*. Generally, therefore, the expression pattern of a chimeric protein will reflect that of the native protein. In cases where more than one virulence factor acts in a complex, however, expressing them individually may not be predictive of their behaviour during a natural infection. Further, the actions of some virulence factors may be antagonistic, as seen for the effect of DcEntA and DcEntB on AMP gene expression, so alone their effects will not reproduce the natural pathophysiology of infection. Nevertheless, when, as here, mutants are not available, heterologous expression can provide one route to understanding virulence factor function under conditions that are more physiological than *ex vivo* or *in vitro* systems.

Dozens of the *D. coniospora* proteins predicted to be secreted are lineage-specific and presumable result from co-evolution with nematode hosts. They are of great interest for the understanding of evolutionary dynamics, for which *D. coniospora* is a potentially powerful model (Courtine et al., 2020), but represent a major challenge due to the lack of any prior knowledge. Therefore, here, we chose to focus on proteins that on the basis of conserved domains were expected to play a direct role in fungal virulence. Thus the first candidate, g2698/RJ55_02698, contains a RIP domain, known to inactivate host ribosomes (Walsh et al., 2013). Given the highly pleiotropic consequences that resulted from its expression, consistent with a non-specific shutdown of protein synthesis, we did not pursue its study. It could, however, provide a good model for blocking ribosome function in *C. elegans*, especially given the possibility to modulate its expression using a combination of RNAi and the auxin inducible degron system.

The three candidate enterotoxin genes that we selected from the expanded genomic repertoire of *D. coniospora* share a highly conserved enterotoxin α domain. Expression of one of them, DcEntC, despite being at a higher level than the other two, was not associated with strong phenotypes, and did not reduce the lifespan of *C. elegans*. Although surprising, this indicates that the effects of DcEntA and DcEntB, which made worms sick and die precociously are specific. They presumably reflect the functions of their auxiliary protein domains.

Worms expressing DcEntA became Dpy as they aged and had a more fragile cuticle. A well-characterised cause of Dpy phenotypes is mutations that affect cuticle collagens, including loss of function mutations in *dpy-7* (Johnstone et al., 1992) and *dpy-10* (Levy et al., 1993), which can also be associated with cuticle fragility (Fechner et al., 2018). This phenotype is shared with *dapk-1* (Tong et al., 2009) and some *bus* mutants (Gravato-Nobre et al., 2005). In all cases, such mutants exhibit an elevated constitutive expression of the p38 MAPK-dependent *nlp* genes (Dodd et al., 2018; Pujol et al., 2008b; Tong et al., 2009; Zugasti et al., 2016). Unexpectedly, DcEntA did not induce, but rather strongly inhibited AMP gene expression. Blocking key immune defence pathways is a strategy used by many pathogens across kingdoms (Lo Presti et al., 2015; Reddick and Alto, 2014; Thakur et al., 2019). Thus, in common with other known virulence factors, for example the *Shigella flexneri* protein effector OspF (Arbibe et al., 2007), DcEntA blocks the activity of a p38 MAPK pathway, in this case with an action downstream of PMK-1. We observed that DcEntA disrupts the normal vesicular pattern of SNF-12. Upon infection, SNF-12 plays an essential role in the induction of AMPs, through its interaction with STA-2 (Dierking et al., 2011). We recently demonstrated that upon wounding, SNF-12 is recruited to the injury site, in a microtubule dependent manner (Taffoni et al., 2020) and that this is crucial for AMP gene expression. Thus DcEntA may inhibit AMP gene expression indirectly through an effect on SNF-12 localisation and activity. This could be sufficient to explain the observed inhibition of STA-2 nuclear translocation, although the results from our mass spectrometry analysis with DcEntA hinted also at a possible interaction with components of the nuclear pore complex, required for the import of proteins into the nucleus.

Suppressing STA-2 activity had the unexpected consequence of promoting the expression of the uncharacterized nematode-specific gene *F40H7.12*, which is highly induced upon *D. coniospora* infection. This could be seen as a fail-safe surveillance mechanism, whereby fungal interference with a major defence pathway leads to a boost of a complementary defence mechanism. The pathway that positively regulates *F40H7.12* is currently unknown. Like *irg-1*, *F40H7.12* expression is induced by translation inhibition. On the other hand, *irg-1* is expressed in the intestine, under the control of the bZIP transcription factor ZIP-2, which is not expressed in the epidermis (Dunbar et al., 2012; Estes et al., 2010). Many enterotoxin α proteins ADP-ribosylate elongation factor family (EF2) proteins essential for ribosome function. Notably, Exotoxin A from *P. aeruginosa* targets EF2, thereby blocking protein synthesis in *C. elegans* intestinal epithelial cells (Dunbar et al., 2012; McEwan et al., 2012). It is possible that DcEntA also acts at the translational level to prevent host defence protein expression as, while not finding EF2 proteins, we noted several eukaryotic Initiation Factor (eIF) proteins, indispensable for translation, among the specific interactors of DcEntA. Together our results indicate that DcEntA affects, directly and indirectly, host defence protein expression, by targeting different cellular processes. This is not unusual for virulence factors, with, for example, EspF from enteropathogenic and enterohemorrhagic *Escherichia coli* described as a “bacterial pathogen's Swiss army knife” because of the diversity of its actions (Holmes et al., 2010). Further study will be required to validate the many candidate host protein interactors and to determine whether any have preponderant roles in pathogenesis during a normal infection.

The effect of DcEntB expression was similarly complex. One prominent consequence was an increase in AMP gene expression, dependent on the canonical STA-2 pathway. DcEntB was concentrated in the nucleolus, a dynamic sub-nuclear organelle for ribosomal RNA (rRNA) biogenesis that acts as a cellular stress sensor (Olson, 2004). For example, impairment of nucleolar function is thought to stabilize p53, a key regulator of cellular homeostasis (Rubbi and Milner, 2003), while inhibition of proteasome activity leads to sequestration of p53 proteins to the nucleolus (Klibanov et al., 2001). In *C. elegans,* the p53 homologue, CEP-1, acts downstream of NOL-6, a nucleolar RNA-associated protein (NRAP), via its transcriptional target SYM-1 to enhance resistance to bacterial infection (Fuhrman et al., 2009). More recently, it was shown that infection by *P. aeruginosa* decreases the level of the nucleolar pre-rRNA processing protein fibrillarin, FIB-1 (Tiku et al., 2018). FIB-1 acts downstream of the homologue BRAT/TRIM2, NCL-1, to regulate rRNA abundance and nucleolar size (Yi et al., 2015). Bacterial infection therefore decreases rRNA and nucleolar size (Tiku et al., 2018).

For the moment, no *P. aeruginosa* effector has been identified that localises specifically to the host nucleolus. Indeed, while it has been known for many years that different RNA-virus proteins traffic to the nucleolus and recruit nucleolar proteins to facilitate viral replication (Hiscox, 2007), it was only comparatively recently that examples of bacterial effectors that target the nucleolus were identified, the first being EspF (Dean et al., 2010). Here, we found that DcEntB is recruited to nucleoli and can make them larger and irregularly shaped. This is also one of the consequences of infection of the epidermis by *D. coniospora*, but whether this depends solely on the action of DcEntB remains to be established. The relationship between the change in nucleolar morphology and the expression of AMP genes appears complex. On the one hand, blocking protein translation, which alters the nucleolus, was associated with a small and specific increase in *nlp* AMP gene expression. On the other, blocking the elevated AMP gene expression induced by DcEntB by knocking down *sta-2* expression reverted nucleoli almost to normal. Further work will be needed to tease out the underlying causal links.

Regardless, the induction of AMPs could be interpreted as a type of surveillance immunity. Potentially, the changes in cellular physiology provoked by DcEntB could be detected as a damage signal and induce an immune response in the epidermis (Figure 11). Several other examples illustrate the important role of surveillance immunity in *C. elegans* (reviewed in (Pukkila-Worley, 2016)). In one case, Stx1, a virulence factor from enterohemorrhagic *E. coli* that is able to inhibit protein synthesis, activates the intestinal p38 MAPK pathway (Chou et al., 2013). Apart from the core MAPK cassette that is shared between epidermis and intestine, the p38 pathway has distinct inputs and outputs in the two tissues (reviewed in (Kim and Ewbank, 2018)). Whether DcEntB exerts similar effects in the intestine is currently not known. It has been shown, however, that there is an intimate balance of MAPK activity between the 2 tissues, with stimulation of the p38 pathway in one negatively influencing its activity in the other (Zugasti et al., 2016), via a mechanism potentially involving the Tribbles homologue NIPI-3 (Kim et al., 2016; McEwan et al., 2016). Thus during the course of an infection, beyond the cell autonomous effects described here, virulence factors like DcEntA and DcEntB have the potential to influence immune defences in distant tissues.

It is interesting to note that in contrast to DcEntA that has a C-terminal domain not found in any other species, DcEntB has orthologues in many pathogenic fungi. These include in nematode-trapping fungi *Dactylellina* spp. (Meerupati et al., 2013) and *Drechslerella* spp. (Liu et al., 2014), as well as the egg-infecting species *Pochonia chlamydosporia* (Larriba et al., 2014). The different species represent distinct branches on the phylogenetic tree, reflecting the multiple independent origins of nematode parasitism (Lebrigand et al., 2016). Orthologues are also found in ant-infecting *Ophiocordyceps* species (de Bekker et al., 2017). It has previously been suggested that heat-labile enterotoxins are important effectors in host adaptation and co-evolution (Kobmoo et al., 2018). It is possible that as a more ancient virulence factor, *C. elegans* has been able to develop a counter-defensive strategy against DcEntB. This would then potentially drive enterotoxin diversification in *D. coniospora*, leading to the emergence of DcEntA. We hypothesise that *C. elegans* has not yet evolved an effective defence strategy against this more recent virulence factor. It will clearly be important in the future to assay the expression of the different enterotoxin genes, and measure the levels of the corresponding proteins in *C. elegans* during an infection, to gauge their relative importance in pathogenesis, which is beyond the scope of the current study.

In conclusion, through this initial investigation of *D. coniospora* virulence factors, we have revealed the very complicated, sometimes antagonistic, nature of some of the molecular interactions that come into play during natural fungal infection of *C. elegans*. In addition to providing insight into the molecular function of two representative enterotoxins, we have gained a new understanding of host defence mechanisms.

## SUPPLEMENTARY FIGURE LEGENDS

**Figure S1.**
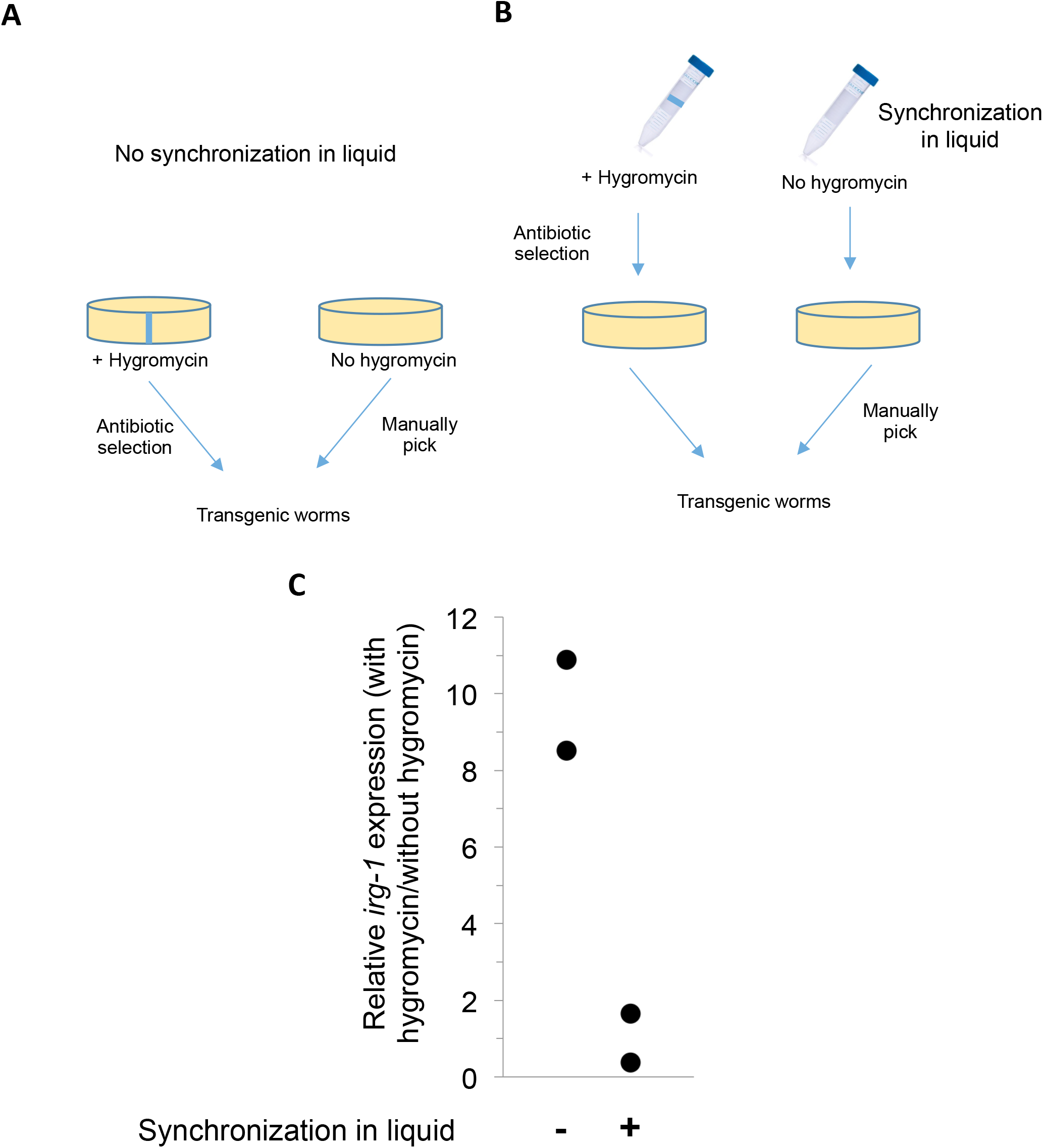
(**A**, **B**) Schematic representation of different culture and selection procedures. (**A**) Transgenic worms carrying *rps-0p::hygR* conferring hygromycin resistance together with *unc-122p::GFP* as an extrachromosomal array (IG1864) were grown on NGM plates supplemented with hygromycin (left) or on standard NGM plates after manual selection on the basis of the expression of the fluorescent marker. (**B**) IG1864 worms were cultured overnight liquid in the presence (left) or absence of hygromycin. Worms were transferred to NGM plates and in the latter case selected manually, as above. (**C**) Quantitative RT-PCR analysis comparing the expression of *irg-1* in IG1864 worms selected by growth on hygromycin-supplemented NGM plates to that in worms selected manually (−), and in IG1864 worms selected by synchronization in the presence of hygromycin to that in worms selected manually following synchronization in the absence of hygromycin (+). The results from 2 independent experiments are shown. It can be seen that even in worms that are resistant to hygromycin, prolonged culture in the presence of the antibiotic increases *irg-1* expression, while overnight exposure during early development does not.

**Figure S2.**
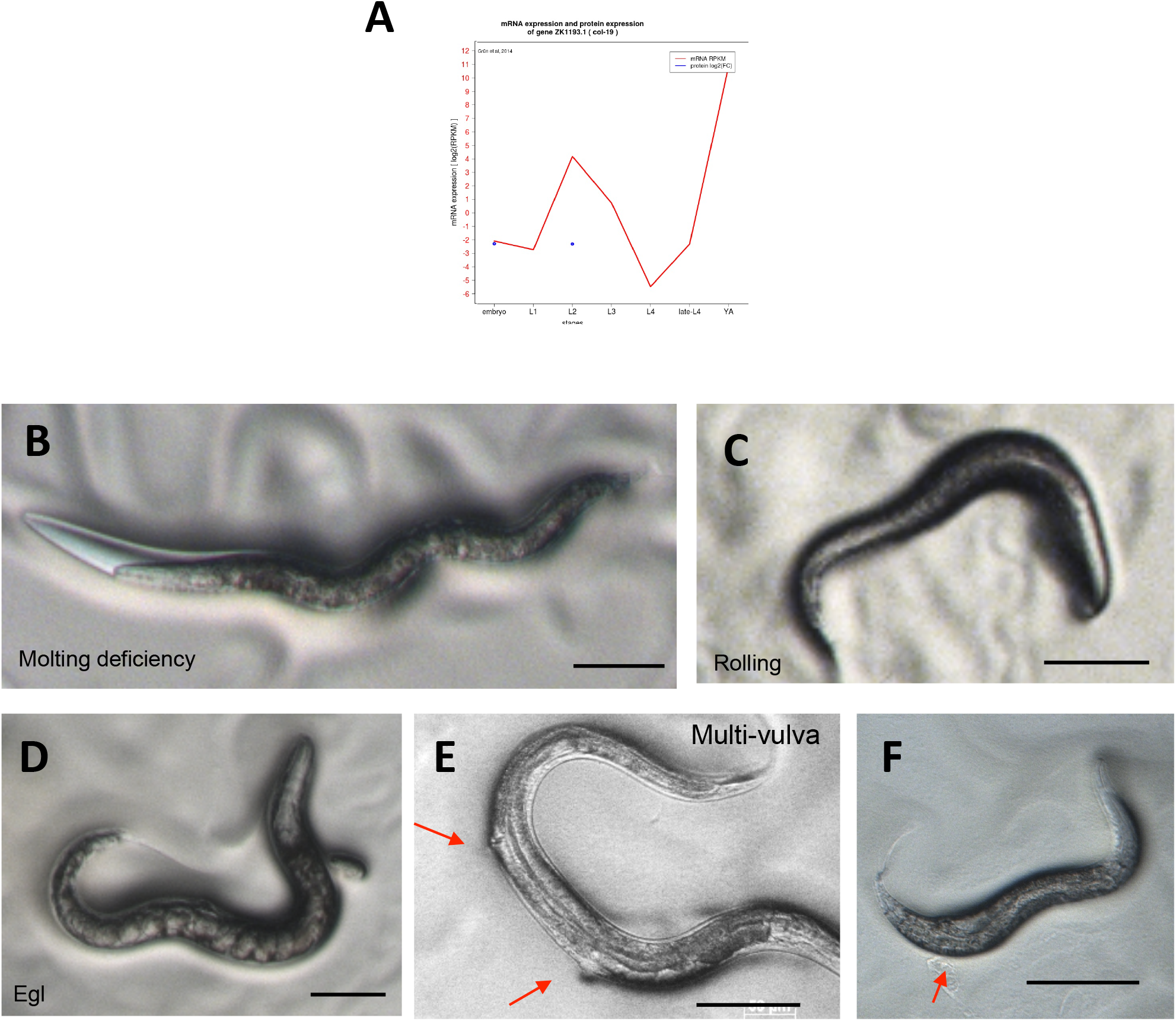
(**A**) Level of expression of *col-19* at different developmental stages. Image adapted from https://elegans.mdc-berlin.de/cel_ex.html. (**B-F**) Photomicrographs of representative worms expressing g2698/RJ55_02698 (IG1867) under the control of the *col-19* promoter and exhibiting moulting defects (**B**), rolling (**C**), egg retention (**D**), multivulva (arrows, **E**) and morphological defect (highlighted by arrow, **F**). Scale bar 100 μm.

**Figure S3.**
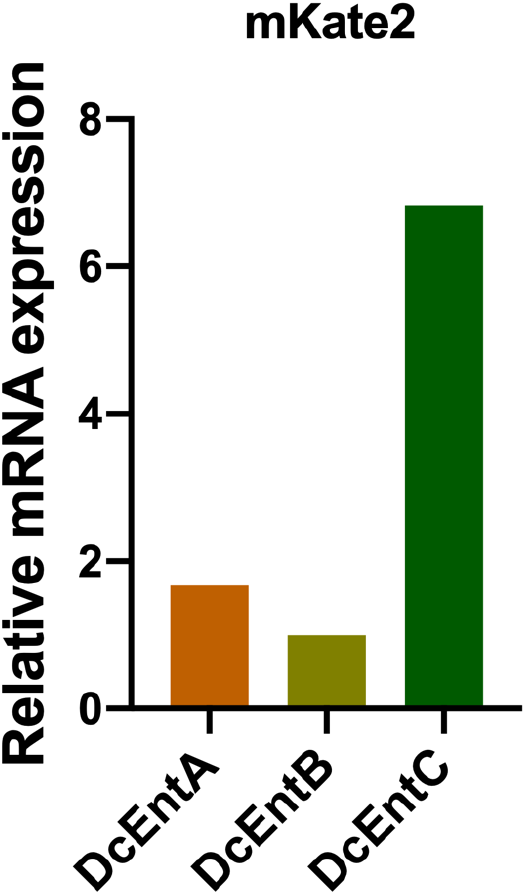
Quantitative RT-PCR analysis of the relative expression of mKate2 in worms expressing DcEntA (IG1926), or DcEntC (IG1880), relative to those expressing DcEntB (IG1925).

**Figure S4.**
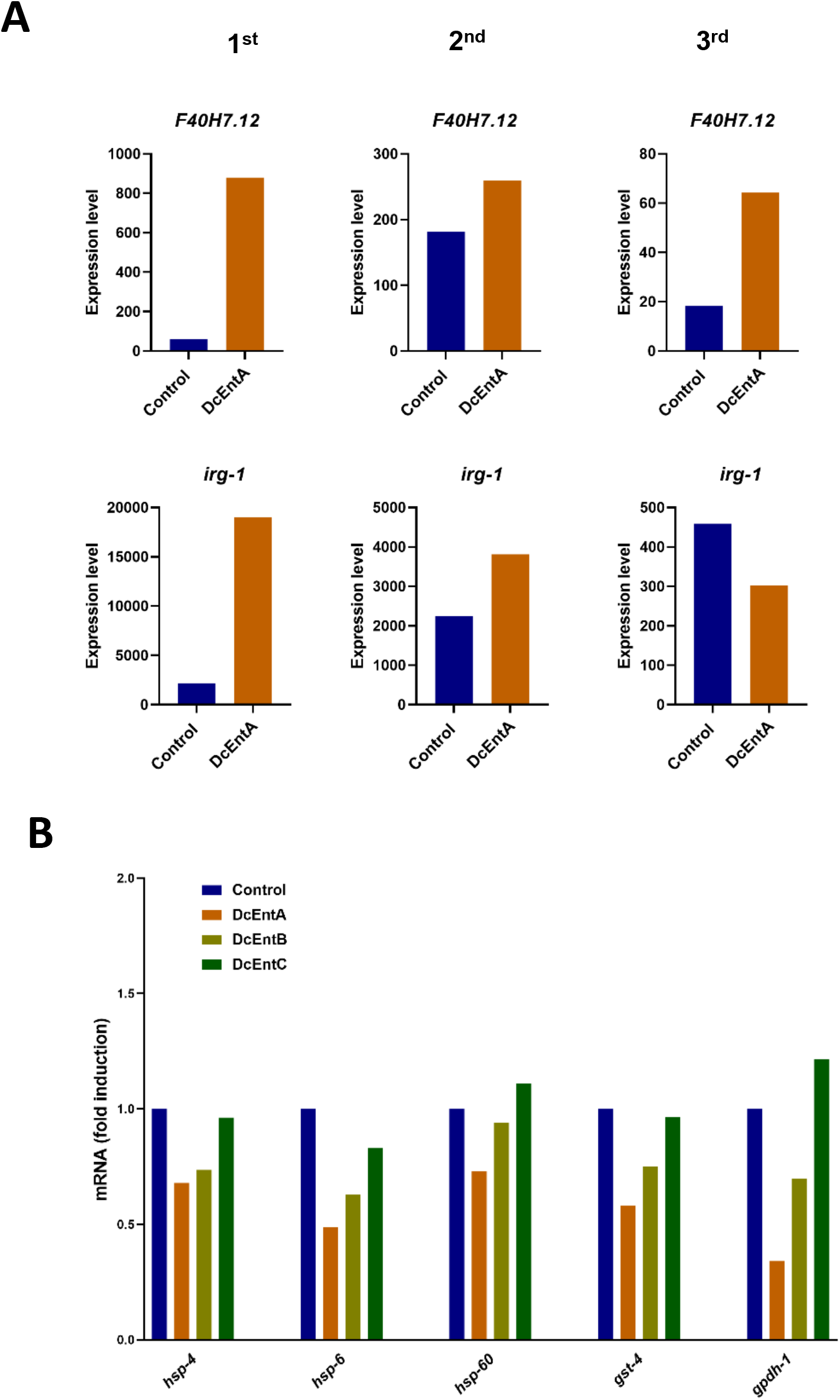
(**A**) Quantitative RT-PCR analysis of the expression of *F40H7.12* and *irg-1* in worms carrying *hygR* and *frIs7* with (IG1942) or without DcEntA (control; IG1864). Data from three independent experiments are shown. (**B**) Quantitative RT-PCR analysis of the expression of *hsp-4*, *hsp-6*, *hsp-60*, *gst-4* and *gpdh-1* in worms expressing DcEntA (IG1926), DcEntB (IG1925) or DcEntC (IG1880). Results are presented relative to control worms (JDW141).

**Figure S5.**
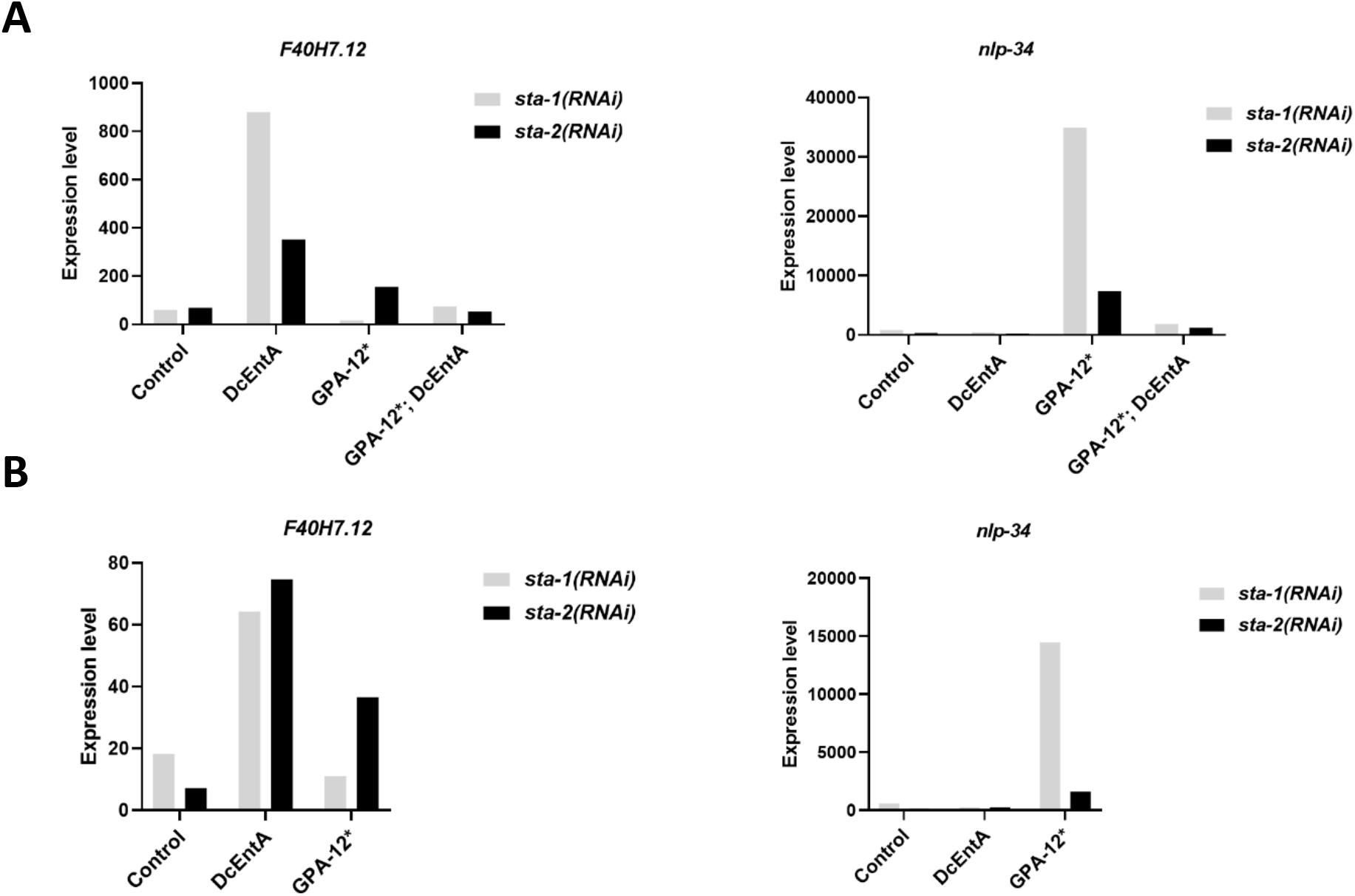
(**A**, **B**) Quantitative RT-PCR analysis of the expression of *F40H7.12* and *nlp-34* in control (*hygR;frIs7* IG1864) worms and worms expressing DcEntA (IG1942), GPA-12* (IG1389), and GPA-12*;DcEntA (IG1948) following RNAi against *sta-1* or *sta-2*.

**Figure S6.**
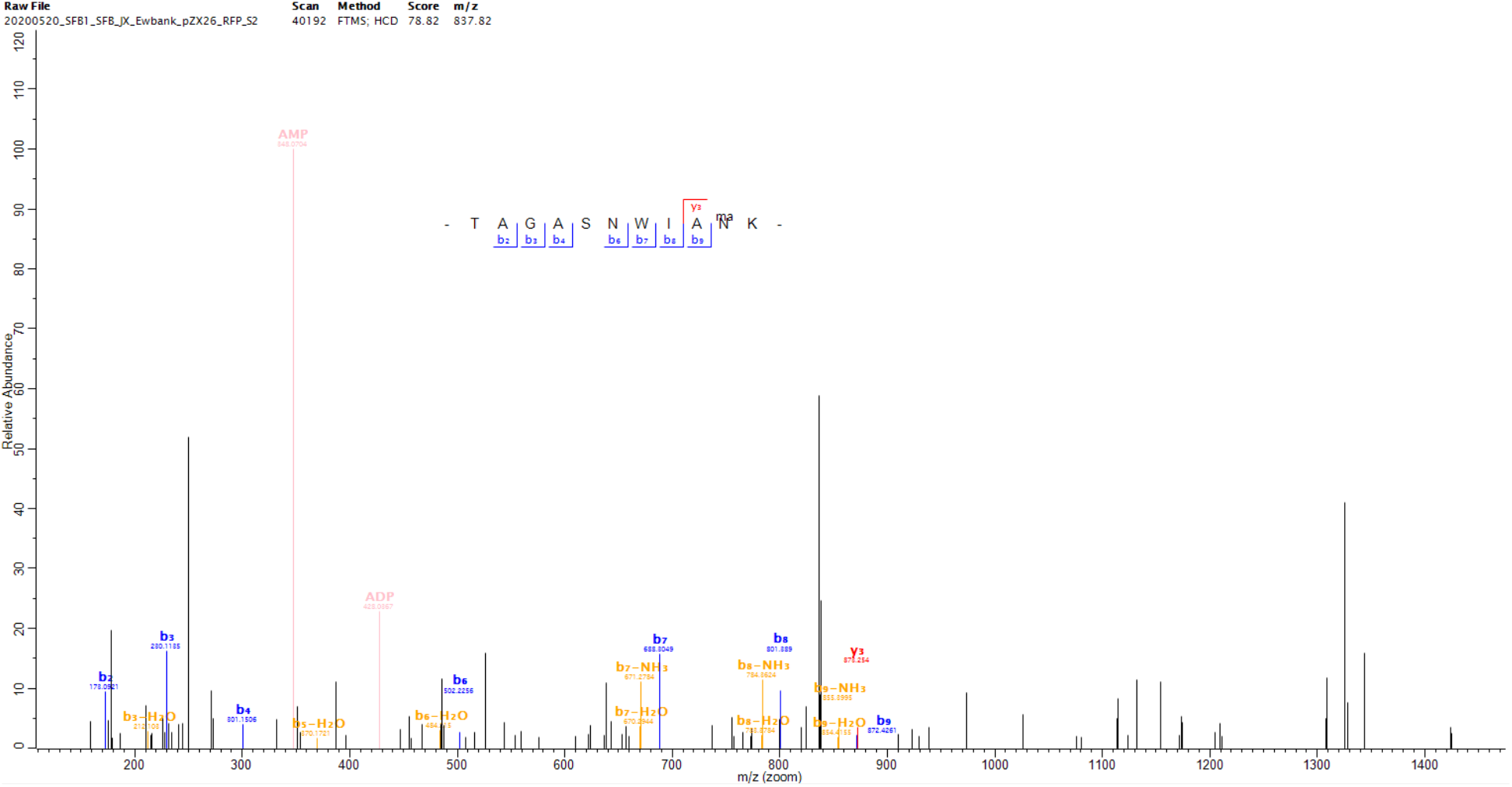
HCD MS/MS spectrum of peptide sequence TAGASNWIANK, identifying asparagine N275 as mono-ADP-ribosylation (MAR) site of DcEntA. Diagnostic ions of AMP and ADP were generated by the ADP-ribosylation group of the modified peptide during HCD fragmentation.

**Figure S7.**
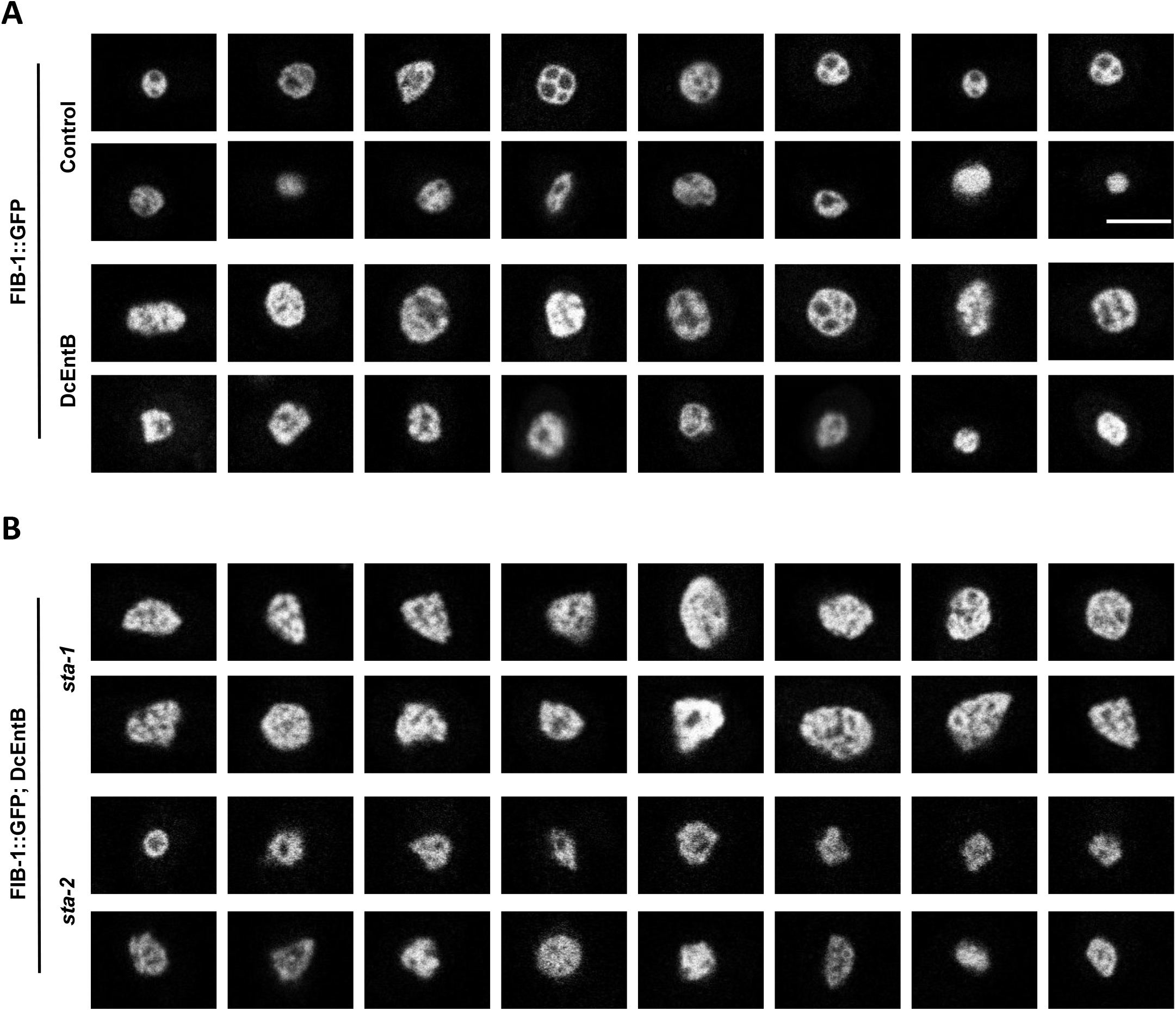
(**A**) Representative confocal images of hyp7 nuclei in young adult worms expressing FIB-1::GFP with (DcEntB, IG1984; lower two panels) or without (Control; SJL1; upper two panels) DcEntB, scale bar, 5 μm. (**B**) Representative confocal images of hyp7 nuclei in young adult worms expressing FIB-1::GFP; DcEntB (IG1984) on *sta-1* (upper two panels) or *sta-2* (lower two panels) RNAi. Scale bar, 5 μm.

**Figure S8.**
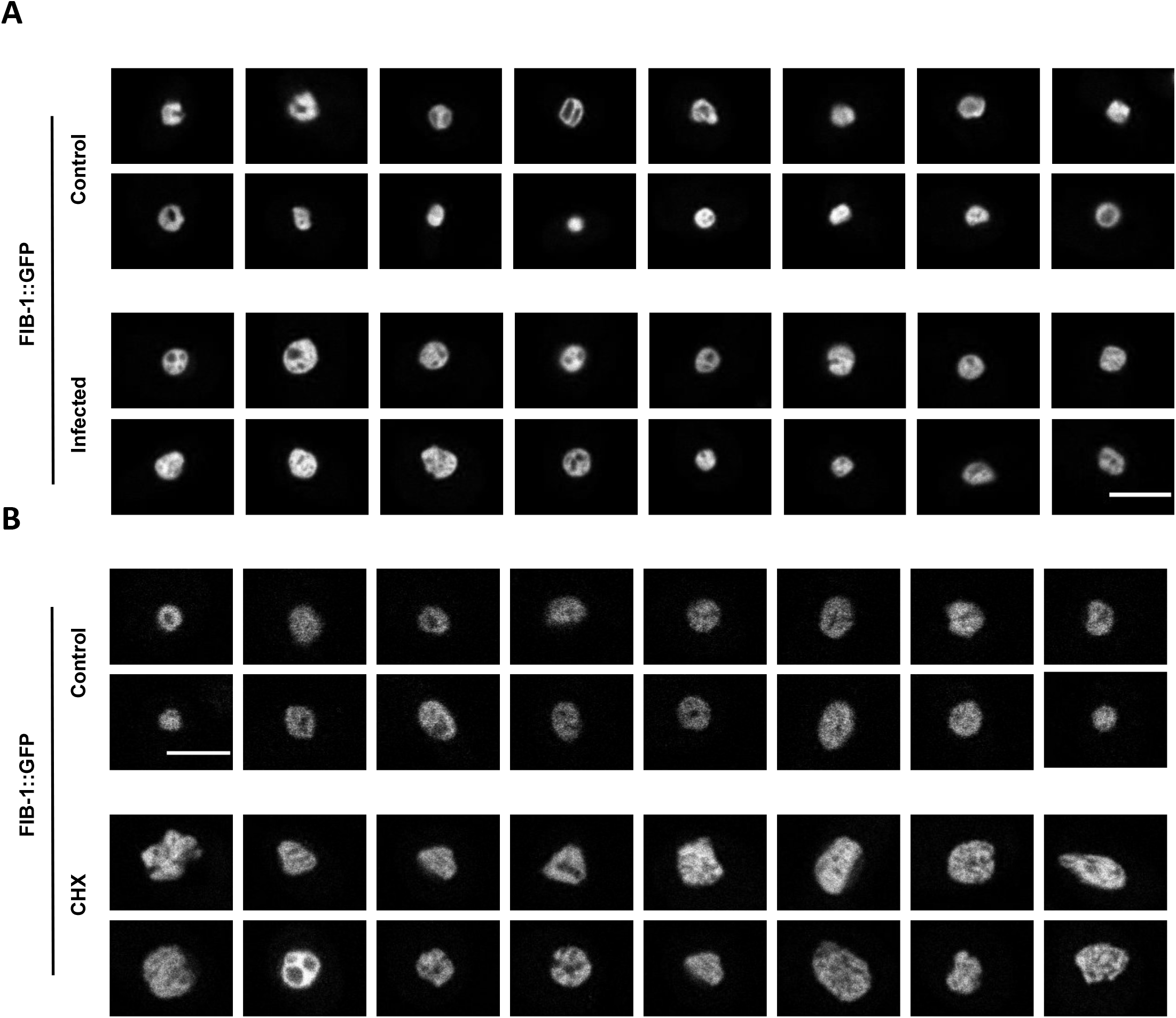
(**A**) Representative confocal images of hyp7 nuclei in young adult SJL1 worms expressing FIB-1::GFP after 24 h infection as young adults at 25°C (Infected; lower two panels) or in uninfected animals (Control; upper two panels). Scale bar, 5 μm. (**B**) Representative confocal images of hyp7 nuclei in young adult SJL1 worms expressing FIB-1::GFP after 6 h CHX exposure (CHX; lower two panels) or without CHX exposure (Control; upper two panels). Scale bar, 5 μm.

## SUPPLEMENTARY TABLES

**Supplementary Table S1. Full genotypes of transgenic strains.**

**Supplementary Table S2. Oligonucleotide primers.**

**Supplementary Table S3. Analysis of predicted secreted *D. coniospora* proteins.** The 225 proteins previously lacking homologues in Genbank (Lebrigand et al., 2016) were analysed.

**Supplementary Table 4. Phenotypes of transgenic worms expressing one of the 3 candidate virulence factors.**

**Supplementary Table 5. Identification of protein-protein interactors for DcEntA.** The first sheet gives quantitative summary statistics for the significant and specific candidate protein-protein interactors for DcEntA obtained from results for analyses of 3 independent samples, referenced to Wormbase release WS275. The subsequent sheets report annotations and gene enrichments using Wormbase tools and WS277. The GeneIDs of the candidate interactors did not evolve between WS275 and WS277.

**Supplementary Table 6. Identification of protein-protein interactors for DcEntB.** The first sheet gives quantitative summary statistics for the significant and specific candidate protein-protein interactors for DcEntB obtained from results for analyses of 3 independent samples, referenced to Wormbase release WS275. The subsequent sheets report annotations and gene enrichments using Wormbase tools and WS277. The GeneIDs of the candidate interactors did not evolve between WS275 and WS277.

## ACKNOWLEDGMENTS

Supported by institutional grants from the Institut national de la santé et de la recherche médicale, Centre national de la recherche scientifique and Aix-Marseille University to the CIML, and the Agence Nationale de la Recherche program grants (ANR-16-CE15-0001-01, ANR-11-LABX-0054 (Labex INFORM) and ANR-11-IDEX-0001-02 (A*MIDEX)). ZX was supported by a fellowship from the China Scholarship Council, DA by the Labex INFORM, and DC by the Turing Centre (ANR-16-CONV-0001; A∗MIDEX). Worm sorting was performed by Jerome Belougne using the facilities of the French National Functional Genomics platform, supported by the GIS IBiSA and Labex INFORM. We thank the imaging core facility (ImagImm) of the Centre d'Immunologie de Marseille-Luminy (CIML), supported by the French National Research Agency program (France-BioImaging; ANR-10-INBS-04-01). We thank Shizue Omi for her contribution, Pierre Golstein for constructive criticism, and Andrew Chisholm for reagents. Some strains were provided by the *Caenorhabditis* Genetics Center (CGC) that is supported by the National Institutes of Health - Office of Research Infrastructure Programs (P40 OD010440). Others were kind gifts from Adam Antebi, Xiaochen Wang, Jordan Ward and Benjamin Weaver.

