## Supplementary Table S1 for "Antagonistic fungal enterotoxins intersect at multiple levels with host innate immune defences"

**Table S1. Full genotypes of transgenic strains.**

| Strain Name | Genotype | Reference |
|---|---|---|
| IG274 | *wt; frIs7[nlp-29p::GFP, col-12p::DsRed] x3* | (Pujol et al., 2008) |
| IG1389 | *wt; frIs7[nlp-29p::GFP, col-12p::DsRed] IV; frIs30[(col-19p::GPA-12gf), pNP21(pBunc-53::GFP)] I* | (Labed et al., 2012) |
| IG1864 | *frIs7[nlp-29p::GFP, col-12p::DsRed]IV; frEx613[unc-122p:GFP, rps-0p::HygR]* | This study |
| JDW141 | *Si[eft-3p::TIR::P2A::BFP-NLS-degron:_3'tbb-2 ] I* | (Ashley et al., 2020) |
| IG823 | *wt; frIs43[col-12p::SNF-12::GFP, ttx-3p::DsRed2] V* | (Dierking et al., 2011) |
| XW18234 | *qxIs727(sta-2p::sfGFP::STA-2, single-copy)* | (Miao et al., 2020) |
| BPW24 | *pmk-1(miy[PMK-1(D327E)]; frIs7 [nlp-29p::GFP, col-12p::DsRed] IV* | (Weaver et al., 2020) |
| SJL1 | *cguIs1[fib-1p::FIB-1::GFP::fib-1 3'UTR]* | (Yi et al., 2015) |
| PX627 | *fxIs1([pie-1p::TIR1::mRuby, I:2851009]) I; spe-44(fx110[spe-44::degron]) IV* | (Kasimatis et al., 2018) |
| IG1502 | *rde-1(ne219) V; Is[wrt-2p::RDE-1_3'unc-54, myo-2p::RFP3]; frIs7[nlp-29p::GFP, col-12p::DsRed IV]* | (Zugasti et al., 2014) |
| IG1867 | *Si[eft-3p::TIR::P2A::BFP-NLS-degron:_3'tbb-2] I; frEx612[pZX21(col-19p::g2698::FLAG::Degron::mKate_3'unc-54), unc-122p:GFP, rps-0p::HygR]* | This study |
| IG1880 | *Si[eft-3p::TIR::P2A::BFP-NLS-degron:_3'tbb-2] I; frEx614[pZX19(col-19p::g6833::FLAG::Degron::mKate_3'unc-54), unc-122p:GFP, rps-0p::HygR]* | This study |
| IG1925 | *Si[eft-3p::TIR::P2A::BFP-NLS-degron:_3'tbb-2] I; frEx619[pZX25(col-19p::g2819::FLAG::Degron::mKate_3'unc-54), unc-122p:GFP, rps-0p::HygR]* | This study |
| IG1926 | *Si[eft-3p::TIR::P2A::BFP-NLS-degron:_3'tbb-2] I; frEx620[pZX26(col-19p::g7949::FLAG::Degron::mKate_3'unc-54), unc-122p:GFP, rps-0p::HygR]* | This study |
| IG1883 | *frEx614[pZX19(col-19p::g6833::FLAG::Degron::mKate_3'unc-54), unc-122p:GFP, rps-0p::HygR]; frIs7[nlp-29p::GFP, col-12p::DsRed] IV* | This study |
| IG1941 | *frEx619[pZX25(col-19p::g2819::FLAG::Degron::mKate_3'unc-54), unc-122p:GFP, rps-0p::HygR]; frIs7[nlp-29p::GFP, col-12p::DsRed] IV* | This study |
| IG1942 | *frEx620[pZX26(col-19p::g7949::FLAG::Degron::mKate_3'unc-54), unc-122p:GFP, rps-0p::HygR]; frIs7[nlp-29p::GFP, col-12p::DsRed] IV* | This study |
| IG1948 | *frEx620[pZX26(col-19p::g7949::FLAG::Degron::mKate_3'unc-54), unc-122p:GFP, rps-0p::HygR]; frIs7[nlp-29p::GFP, col-12p::DsRed] IV; frIs30[(col-19p::GPA-12gf), pNP21(pBunc-53::GFP)] I x3* | This study |
| IG1963 | *frEx620[pZX26(col-19p::g7949::FLAG::Degron::mKate_3'unc-54), unc-122p:GFP, rps-0p::HygR];pmk-1(miy[PMK-1(D327E)];frIs7[nlp-29p::GFP, col-12p::DsRed] IV* | This study |
| IG1971 | *qxIs727(sta-2psfGFP::STA-2,single-copy); frEx620[pZX26(col-19p::g7949::FLAG::Degron::mKate_3'unc-54), unc-122p:GFP, rps-0p::HygR]; Si[eft-3p::TIR::P2A::BFP-NLS-degron:_3'tbb-2] I* | This study |
| IG1977 | *qxIs727(sta-2psfGFP::STA-2,single-copy); frEx619[pZX25(col-19p::g2819::FLAG::Degron::mKate_3'unc-54), unc-122p:GFP, rps-0p::HygR]; Si[eft-3p::TIR::P2A::BFP-NLS-degron:_3'tbb-2] I* | This study |

| | | |
|---|---|---|
| IG1984 | *Si[eft-3p::TIR::P2A::BFP-NLS-degron:_3'tbb-2] I; frEx619[pZX25(col-19p::g2819::FLAG::Degron::mKate_3'unc-54), unc-122p:GFP, rps-0p::HygR] ; cguIs1[fib-1p::FIB-1::GFP::fib-1 3'UTR]* | This study |
| IG1998 | *frIs43[col-12p::SNF-12::GFP, ttx-3p::DsRed2] V; frEx620[pZX26(col-19p::g7949::FLAG::Degron::mKate_3'unc-54), unc-122p:GFP, rps-0p::HygR]* | This study |


Ashley, G., T. Duong, M.T. Levenson, M.A.Q. Martinez, J.D. Hibshman, H.N. Saeger, R. Doonan, N.J. Palmisano, R. Martinez-Mendez, B. Davidson, W. Zhang, J.M. Ragle, T.N. Medwig-Kinney, S.S. Sirota, B. Goldstein, D.Q. Matus, D.J. Dickinson, D.J. Reiner, and J.D. Ward. 2020. Expanding the *Caenorhabditis elegans* auxin-inducible degron system toolkit with internal expression and degradation controls and improved modular constructs for CRISPR/Cas9-mediated genome editing. *bioRxiv*:2020.2005.2012.090217.

Dierking, K., J. Polanowska, S. Omi, I. Engelmann, M. Gut, F. Lembo, J.J. Ewbank, and N. Pujol. 2011. Unusual regulation of a STAT protein by an SLC6 family transporter in *C. elegans* epidermal innate immunity. *Cell Host Microbe*. 9:425-435.

Kasimatis, K.R., M.J. Moerdyk-Schauwecker, and P.C. Phillips. 2018. Auxin-Mediated Sterility Induction System for Longevity and Mating Studies in *Caenorhabditis elegans*. *G3 (Bethesda)*. 8:2655-2662.

Labed, S.A., S. Omi, M. Gut, J.J. Ewbank, and N. Pujol. 2012. The pseudokinase NIPI-4 is a novel regulator of antimicrobial peptide gene expression. *PLoS One*. 7:e33887.

Miao, R., M. Li, Q. Zhang, C. Yang, and X. Wang. 2020. An ECM-to-Nucleus Signaling Pathway Activates Lysosomes for *C. elegans* Larval Development. *Dev Cell*. 52:21-37 e25.

Pujol, N., S. Cypowyj, K. Ziegler, A. Millet, A. Astrain, A. Goncharov, Y. Jin, A.D. Chisholm, and J.J. Ewbank. 2008. Distinct innate immune responses to infection and wounding in the *C. elegans* epidermis. *Curr Biol*. 18:481-489.

Weaver, B.P., Y.M. Weaver, S. Omi, W. Yuan, J.J. Ewbank, and M. Han. 2020. Non-Canonical Caspase Activity Antagonizes p38 MAPK Stress-Priming Function to Support Development. *Dev. Cell*. 53:358-369.e356.

Yi, Y.H., T.H. Ma, L.W. Lee, P.T. Chiou, P.H. Chen, C.M. Lee, Y.D. Chu, H. Yu, K.C. Hsiung, Y.T. Tsai, C.C. Lee, Y.S. Chang, S.P. Chan, B.C. Tan, and S.J. Lo. 2015. A Genetic Cascade of *let-7-ncl-1-fib-1* Modulates Nucleolar Size and rRNA Pool in *Caenorhabditis elegans*. *PLoS Genet*. 11:e1005580.

Zugasti, O., N. Bose, B. Squiban, J. Belougne, C.L. Kurz, F.C. Schroeder, N. Pujol, and J.J. Ewbank. 2014. Activation of a G protein-coupled receptor by its endogenous ligand triggers the innate immune response of *Caenorhabditis elegans*. *Nat Immunol*. 15:833-838.
