## Supplementary Table S2 for "Antagonistic fungal enterotoxins intersect at multiple levels with host innate immune defences"

### Table S2. Oligonucleotide primers

#### Fusion PCR primers

| JEP | Sequence name | Sequence |
|---|---|---|
| 2247 | pDest_pcol-19 R | gttgatgaactgatgtctttc |
| 2925 | 3'unc-54 | catctcgcgcccgtgcctctgacttc |
| 3232 | g4535-F | gaaagacatcagttcatcaacgcggccgcATGTGTTCCGGAGGCTCCAACG |
| 3233 | g4535-R | TCGTCATCCTTGTAATCatcgatGTAGTCAGGGGATGTTGCTTTGG |
| 3234 | FLAG+MKATE-F | atcgatGATTACAAGGATGACG |
| 3235 | FLAG+MKATE-R | gaggcacgggcgcgagatgTTAACGGTGTCCGAGCTTGGATGGGAGG |
| 3236 | prps-0:HygR-F | cactatagggcgaattgggtaccATTTTTGCTTTCGTCGTAAATC |
| 3237 | prps-0:HygR-R | atctgatgacagcggccgcggCTATTCCTTTGCCCTCGGACG |
| 3247 | degron-F | AATCCggcgcgccaagcgctgctggttccATGCCTAAAGATCCAGCCAAACC |
| 3248 | degron-R | GGACATggaaccagcagcgcttggcgcgccCTTCACGAACGCCGCCGCC |
| 3251 | g6833-F | gaaagacatcagttcatcaacgcggccgcATGGTTCCGCCACCAGCCTCGCCG |
| 3252 | g6833-R | TCGTCATCCTTGTAATCatcgatACCGCTTGTGTGCCCCTTCTTTGC |
| 3253 | g2698-F | gaaagacatcagttcatcaacgcggccgcATGTTCATTGGCTTTACTGTCACC |
| 3254 | g2698-R | TCGTCATCCTTGTAATCatcgatAGGATGCAACTCGCATTTTCC |
| 3269 | g2819-F | gaaagacatcagttcatcaacgcggccgcATGCACTTGGTCCCATCTGACG |
| 3270 | g2819-R | TCGTCATCCTTGTAATCatcgatGGCCTTTTCGCCGCCAAAACCTTTGC |
| 3271 | g7949-F | gaaagacatcagttcatcaacgcggccgcATGCGCCGGCACAAGCTCAAGCCCG |
| 3272 | g7949-R | TCGTCATCCTTGTAATCatcgatTGAGTCTTTGATTTCCGAGAAGCC |

#### qRT-PCR primers

| JEP | Sequence name | Sequence |
|---|---|---|
| 538 | *act-1* | ccatcatgaagtgcgacattg |
| 539 | *act-1* | catggttgatggggcaagag |
| 952 | *nlp-29* | tatggaagaggatatggaggatatg |
| 848 | *nlp-29* | tccatgtatttactttccccatcc |
| 950 | *nlp-31* | ggtggatatggaagaggttatggag |
| 953 | *nlp-31* | gtctatgcttttactttcccc |
| 969 | *nlp-34* | atatggataccgcccgtacg |
| 970 | *nlp-34* | ctattttccccatccgtatcc |
| 549 | *cnc-2* | tcccatgcccataccgtaac |
| 944 | *cnc-2* | ccgctcaatatggttatggag |
| 1124 | *cnc-4* | acaatggggctacggtccatat |
| 1125 | *cnc-4* | actttccaatgagcattccgagga |
| 1676 | *irg-1* | ccatggaatgaaacttgtgg |
| 1677 | *irg-1* | ccagtttcgttcatcttcaca |
| 2340 | *F40H7.12* | ttcctgagtgctcacgaagg |
| 2341 | *F40H7.12* | aacactgaggaacgaccagg |
| 2863 | *hsp-4* | gccatctcgtggaatcaacc |
| 2864 | *hsp-4* | gtgagtggattgacgtcaag |
| 2876 | *hsp-6* | caacagatcgttatccaatc |
| 2877 | *hsp-6* | ttctctttgcgtcctcagc |
| 3119 | *hsp-60* | gttgaagttggagagaagaaggacc |

| | | |
|---|---|---|
| 3120 | *hsp-60* | atctgagaagagcaacacctcc |
| 3307 | *gst-4* | gaaaatttggactcgctgg |
| 3308 | *gst-4* | aagaaatcatcacgggctgg |
| 1795 | *gpdh-1* | agcactaaagaacattgtcgcc |
| 1796 | *gpdh-1* | tggtaatgagatcagccactcc |
| 3179 | *mKate2* | acaacgtcaagatccgtggagtc |
| 3180 | *mKate2* | ggtaggtggtcttgaggttgc |
