## Supplementary Table S4 for "Antagonistic fungal enterotoxins intersect at multiple levels with host innate immune defences"

| Phenotype | DcEntA | DcEntB | DcEntC |
| --- | --- | --- | --- |
| Low or irregular pumping rate | x | x | x |
| Abnormal locomotion or lethargic | x | | x |
| Protruding vulva | x | x | |
| Bursting (through vulva) | | x | |
| Single gonad arm | x | x | |
| Asymmetric gonad arms | x | | x |
| Egg retention | | | x |
| Major morphological defects (e.g. square head) | x | | |

**Supplementary Table 4. Phenotypes of transgenic worms expressing one of the 3 candidate virulence factors.** Ten worms of each strain cultured at 25˚C were inspected as 2-day old adults. A cross indicates that at least one worm exhibited the phenotype.
